# Extracting Value Coding Features from Individual Serotonin Neurons

**DOI:** 10.1101/2025.07.28.667134

**Authors:** Emerson F. Harkin, Jean-Claude Béïque, Richard Naud

## Abstract

Adaptive behaviour requires animals to continually reevaluate the appetitive or aversive quality of their surroundings. Dorsal raphe serotonin neurons, the main source of serotonergic input to the forebrain, have been implicated in both signaling the quality of an animal’s environment and regulating reward-seeking and punishment-avoiding behaviour, but the precise quantity signaled by these neurons has remained unclear, as well as how these neurons relate with behaviour. Using open-access recordings of serotonergic neurons of the dorsal raphe nucleus while animals perform a dynamic Pavlovian task, we compare firing rate and behavioral data with a model that considers reward history accumulated over a tunable timescale. Our Bayesian parameter estimation supports that serotonergic neurons are consistent with reward history being estimated over about a hundred trials on average, with a heterogeneity across individual neurons spanning 30 to 300 trials. Anticipatory licking also correlated with reward history at multiple timescale, but could not be dissociated from that of a time/thirst nuisance variable and otherwise mostly on a timescale faster than seen in serotonergic cells. These results provide a more precise picture of the dynamics of serotonergic cells under a dynamic Pavlovian task.

## 1 INTRODUCTION

Adaptive behaviour requires animals to continually reevaluate the quality of their environment. The serotonin neurons of the dorsal raphe nucleus (DRN) have been implicated in regulating (1–8) and signaling (9–13) behaviours that relate to environment quality, including fleeing, feeding and mating. Conceptually similar, the concept of state value, central to many reinforcement learning algorithms, is a precisely defined proxy of environment quality. State value has been a central quantity for models of Pavlovian conditioning for decades (14). This quantity was recently shown to predict features of DRN neuron activity, supporting the idea that bulk serotonergic activity in the brain signals state value (12). The precise relationship between single serotonergic neuron activity and the ever changing state value, however, remains unclear as it would require a quantitative model fitting approach. This is challenging because, in reinforcement learning, state value is dynamic, continually updated based on prior experience due to both a reward time horizon (value discounting timescale) and a value estimation timescale. The problem therefore requires that the timescales as well as the relevance of the code be assessed simultaneously. Going further, given that serotonergic and dopaminergic activity follow reward association with different learning timescales (15), the question arises of whether features of the behaviour correlates with value code at the learning timescale found in the majority, only some, or none of the serotonergic neurons. This question can be addressed by fitting anticipatory licking with a model of value evolving over multiple timescales and comparing the timescale of behaviour adaptation with that of the adaptation of responses in serotonergic cells.

Approaches to model fitting and model selection are separated in two broad categories. Given multiple hypotheses, the frequentist approach assigns probability to data and will select the model for which the data is most likely. The Bayesian approach assigns instead a probability to the hypotheses. This is done by building a sufficiently general model and estimating the confidence interval on the parameters that regulate the switch between different hypotheses. Here we used the Bayesian approach to relate a state value model with serotonin neuron activity recorded in mice performing a task with fluctuating reward delivery. We first developed a mathematical framework able to fit a value model on single neuron responses. The result show that different neurons have heterogeneous coding in 1) the degree to which their firing is consistent with state value and 2) the time scale with which the value signal is learned. While different neurons show different learning timescales, the average learning timescale we found was around 100 trials (10-20 min), consistent with a prior report based on population responses (15). Then, we leveraged simultaneous recording of anticipatory licking in the same experiments to ask if this aspect of behaviour was related to value coding at the timescale recorded in the serotonergic neurons. Adapting our Bayesian approach to behaviour, we found that only the value estimated on timescales faster than those associated on average with the value code in the DRN were correlated with anticipatory licking. While some serotonin neurons faster learning timescales than others, they still did not track the evolution of behaviour. Hence our work provides a quantitative report of potency of the value code and its learning timescale in individual serotonin neurons, an observed timescale that generally does not match the timescales correlated with behaviour.

## 2 RESULTS

We used an openly accessible data set where mice performed a dynamic Pavlovian task (Fig. 1; Ref. (16); see Methods). In this task, rewards were associated with a cue based on a probability of reward that changed in piecewise constant blocks. The activity of genetically identified serotonergic cells were recorded using extracellular electrodes. Anticipatory licking was also recorded. While the activity during the randomly initiated trials has previously been shown to relate with state value (12), we focused on the activity in the inter-trial period because the greater number of action potentials makes it more amenable to track learning-related changes and it allows us to ignore the effect of the discounting timescale in a value model since this timescale is shorter than the trial duration (10, 12). This approach is supported by both theory and previous observations, whereby the value of the inter-trial period scales with the value of the trace period (17) (see Discussion).

**Figure 1.**
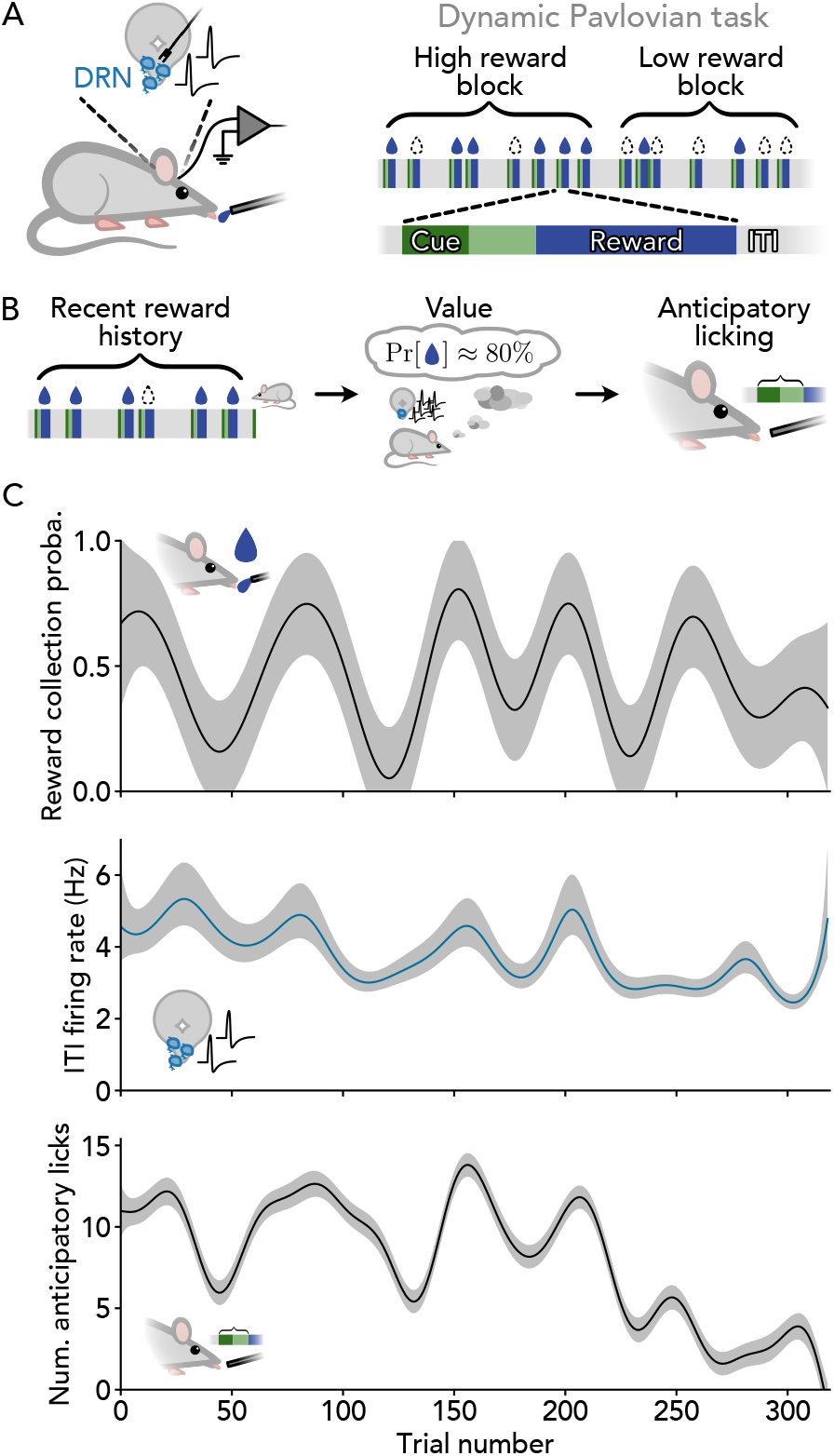
Overview of experimental setup and conceptual model. **A** Experimental setup in the dynamic Pavlovian conditioning task of Grossman et al.(16). In vivo electrophysiological recordings of the activity of individual geneticallyidentifieddorsal raphe nucleus (DRN) serotonin neurons were collected from headfixedmice receiving stochastic water rewards in a trace conditioning task. Trialsconsisted of a 1 s odour cue followed by a 1 s trace period and 3 s reward collectionwindow. Inter-trial interval (ITI) durations followed an exponential distribution with mean 3.3 s. Rewards were delivered with probability 20%, 50%, or 80% in a block structure with random uncued transitions. See Methods and ref. 16 for details. **B** Conceptual model. Recent reward history is used to calculate value, defined as an estimate of the probability of receiving a reward in the upcoming trial. We hypothesize that high value is reflected in both an increase in the ITI firing rates of serotonin neurons and an increase in anticipatory licking. **C** Time-course of reward collection probability (top), serotonin neuron firing rate (middle), and number of anticipatory licks per trial (bottom) for an example recording session. For illustration purposes, all data are smoothed using Gaussian process regression (squared exponential kernel with length scale 20 trials, observation standard deviation set to 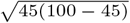 percent, 62ms, and 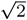 licks for top, middle, and bottom, respectively). Bands represent an approximate 95% CI. Note that firing rates were obtained by analyzing the mean inter-spike interval (ISI) in each ITI, which more closely follows a normal distribution, and inverting the results.

### 2.1 Reward, behaviour, and neural activity evolve slowly over time

As a first step, we sought to determine the timescales over which behaviour and neural activity evolve in this experiment. Frequency analysis of the autocorrelation patterns of reward collection, anticipatory licking, and serotonin neuron firing collectively revealed peaks at two frequencies: one near 1*/*70 cycles trial^*⨪*1^, corresponding to the upper limit of the block length, and another near 1*/*350 cycles trial^*⨪*1^, corresponding to the mean duration of a recording session (Fig. 2). Whereas previous analysis of this data focused on short-term fluctuations over the course of fewer than ten trials (12, 16), these results suggest that both behaviour and neural activity mainly evolve over significantly longer timescales.

**Figure 2.**
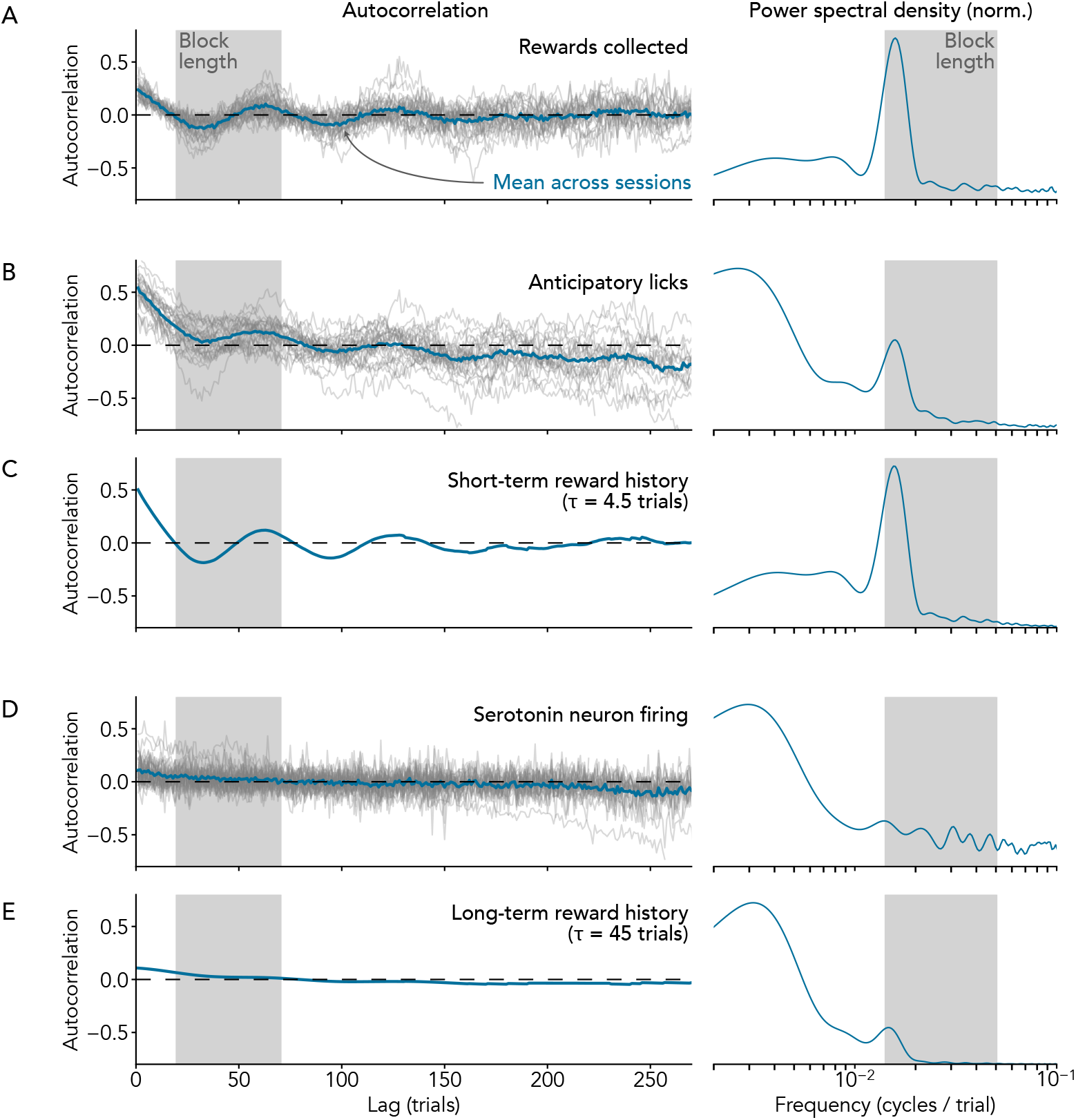
Reward collection, anticipatory licking behaviour, and serotonin neuron firing rates evolve slowly over time in a dynamic Pavlovian task. Left column shows sample autocorrelation (individual sessions or neurons in gray, mean across sessions or neurons in blue), right column shows the estimated power spectrum (computed from the corresponding mean autocorrelation, see Section 5.3.3). **A** Autocorrelation and power spectrum of reward collection. The power spectrum has a peak at 1/63 cycles trial⨪1. N = 28 sessions. **B** Autocorrelation and power spectrum of anticipatory licking. The power spectrum has peaks at 1/377 cycles trial⨪1 and 1/63 cycles trial⨪1. N = 28 sessions. **C** Autocorrelation and power spectrum of an exponentially-weighted moving average of reward collection using a time constant of τ = 4.5 trials. Only the mean across N = 28 sessions is shown. The power spectrum has a peak at 1/64 cycles trial⨪1. Compare with B. **D** Autocorrelation and power spectrum of serotonin neuron firing activity, quantified as the mean inter-spike interval during each inter-trial interval. The power spectrum has a peak at 1/339 cycles trial⨪1. N = 37 neurons. **E** Autocorrelation and power spectrum of an exponentially-weighted moving average of reward collection using a time constant of τ = 45 trials. Only the mean across N = 28 sessions is shown. The power spectrum has a peaks at 1/319 cycles trial⨪1 and 1/69 cycles trial⨪1. Compare with D. Autocorrelations in C and E are normalized such that the autocorrelations at lag one match the means shown in B and D, respectively. For clarity, the autocorrorelation at zero lag is not shown. The 20–70 trial block length used in the experiment and the corresponding frequency range of 1/70–1/20 cycles trial⨪1 are indicated in gray.

Interestingly, fluctuations corresponding to block length and session length were not represented equally in the dynamics of reward collection, anticipatory licking, and serotonin neuron activity. The power spectrum of reward col-lection showed a pronounced peak at 1*/*63 cycles trial^*⨪*1^ and much less power at lower frequencies (Fig. 2A), sug-gesting that the dynamics of reward collection are dominated by the block structure of the experiment rather than session-long trends. In contrast, the power spectrum of an-ticipatory licking showed peaks at both 1*/*63 cycles trial^*⨪*1^ and 1*/*377 cycles trial^*⨪*1^ (Fig. 2B). This pattern is not consistent with the idea that anticipatory licking simply reflects the number of rewards collected in the recent past, since frequency analysis of an exponentially-weighted moving average of collected rewards with a learning timescale of *τ* = 4.5 trials reveals only a single peak corresponding to the block length (Fig. 2C). Finally, analysis of serotonin neuron activity reveals a single peak at 1*/*339 cycles trial^*⨪*1^ (Fig. 2D). This is also not consistent with a short-term history of collected rewards (Fig. 2C), but could arise from a medium-term reward history with a timescale similar to the average block length of 45 trials (Fig. 2E). Overall, we find that the dynamics of reward collection are dominated by fluctuations on a timescale that corresponds to the block length and serotonin neuron activity is dominated by session-long trends, while anticipatory licking displays a mixture of both timescales.

### 2.2 Serotonin neuron activity is consistent with a slowly-evolving estimate of value

To directly examine the connection between serotonin neuron activity and reward history, we constructed a generative model of inter-trial interval firing statistics as a function of value *v*_*t*_

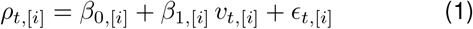

where *ρ*_*t*,[*i*]_ is the firing rate of neuron *i* during inter-trial interval *t*; *β*_0,[*i*]_ is the firing rate intercept, which represents the baseline firing rate when value is zero; *β*_1,[*i*]_ is the firing rate gain, which controls how quickly firing increases or decreases with increasing value; and *ϵ*_*t*,[*i*]_ is a noise term (see Section 5.2.3). We operationally define value as an estimate of the reward probability in the upcoming trial *v*_*t*,[*i*]_ ≈ Pr [*R*_*t*+1,[*i*]_ = 1] learned according to the Rescorla–Wagner rule

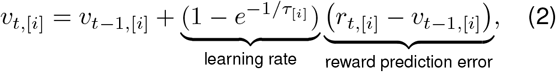

where *r*_*t*,[*i*]_ is the reward received in trial *t, v*_*t*,[*i*]_ is the value during the inter-trial interval immediately following *r*_*t*,[*i*]_, and *τ*_[*i*]_ is the learning timescale. Importantly, value learned in this way is equivalent to an exponentially-weighted moving average of past rewards with learning timescale *τ* and initial condition *v*_0_

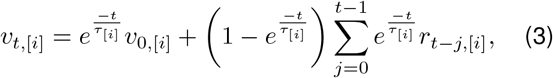

as used in the previous section. Overall, our model assumes that serotonin neuron activity reflects a moving average of recent rewards, where the meaning of “recent” is governed by the learning timescale *τ*. Thus Eq. 2 defines our value model and Eq. 1 defines how it is fit to the observed firing rate (Fig. 3A-B).

**Figure 3.**
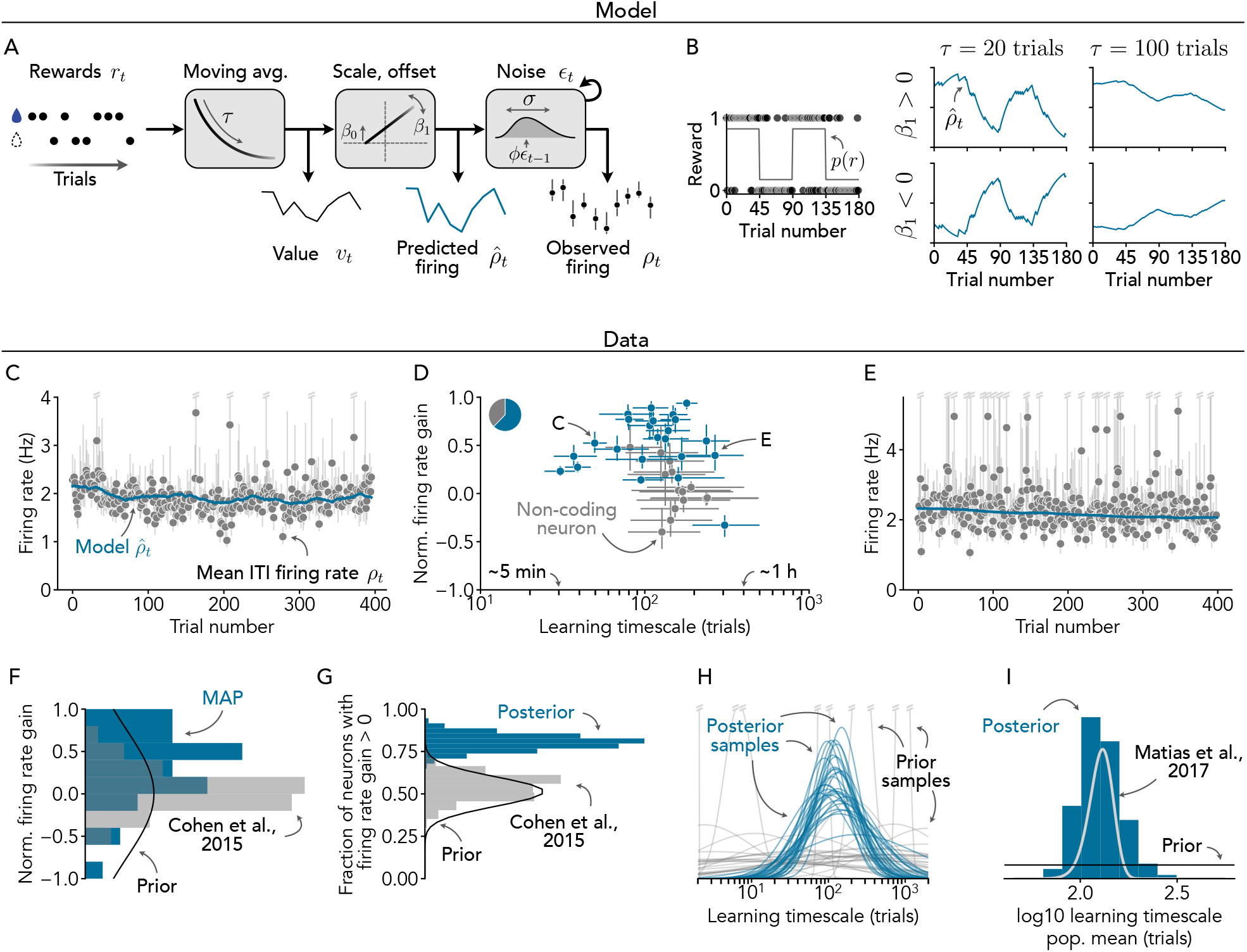
Value coding features of serotonin neurons. **A** Generative model of serotonin neuron activity. Rewards are passed into an exponentially-weighted moving average with timescale τ to estimate value vt (equivalent to learning value according to the Rescorla–Wagner rule) which is then scaled (by gain factor β1) and offset (by intercept β0) to produce an estimated firing rate 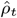. The observed firing rate ρt is a noisy reflection of 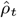. Note that the noise is autocorrelated, causing the observed firing rate to drift. See Methods for details. **B** Effects of varying firing rate gain β1 and learning timescale τ on the predicted firing rate 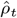. Simulated rewards from alternating 85% and 15% reward probability blocks are shown at left, and model outputs are shown at right. Note that for illustration purposes, here we have used more extreme reward probabilities and more regular block structure than in the actual experiment. **C** Example of a value coding neuron with a short estimated learning timescale. Points and error bars indicate mean and SEM firing rate in each inter-trial interval. Blue line indicates the MAP value model prediction. For clarity, one outlier trial with very high estimated firing rate and the corresponding model prediction are not shown (note small gap in blue line near 375 trials). **D** Estimated normalized firing rate gain and learning timescale for each neuron. Points and error bars represent the median and IQR of the posterior, respectively. Neurons for which the 95% credibility interval for the normalized firing rate gain includes zero are designated putatively non-value-coding and are indicated in gray; the fraction of neurons that neurons that fall in this category (37.8%) is indicated in the pie chart. Exemplar neurons from C and E are labeled. For reference, timescales that correspond to approximately 5 min and 1 h are highlighted based on the average trial plus ITI duration (9.3 s). **E** Example of a value coding neuron with a long estimated learning timescale. Data and model presented as in A. For clarity, two outlier trials with very high estimated firing rate and corresponding model predictions are not shown. **F** Distribution of normalized firing rate gain across neurons. Distribution from the MAP model prediction is shown in blue, and the Bayesian prior used for model fitting is shown in black. For comparison, the distribution of normalized firing rate gain across neurons in ref. 9 is shown in gray. **G** Fraction of N = 37 neurons with positive firing rate gain. Positive firing rate gain indicates that the neuron is activated when value is high. Full posterior distribution is shown in blue, and the Bayesian prior used for model fitting is shown in black. For comparison, the estimated fraction of neurons in the data of ref. 9 are activated when value is high is shown in gray (bootstrap distribution). **H** Distribution of learning timescales in the full population of serotonin neurons. To illustrate uncertainty in this estimate, 30 draws from the posterior are shown. Samples from the prior are shown in gray. **I** Mean learning timescale in the full population of serotonin neurons. Full posterior distribution is shown in blue, and the flat prior is shown in black. For comparison, the estimated mean from ref. 15 is shown in gray. Note close agreement with our results.

To assess whether this value model provides a meaningful account of the activity patterns of serotonin neurons in this experiment, we compared its ability to predict firing in held-out trials against two control models: one which assumes a constant firing rate and the other which assumes a session-wide linear trend. The value model performed significantly better than the two control models in terms of the average error across all neurons in this dataset (mean squared error on the mean inter-spike interval inheld-out trials of 0.0209 s^2^ for the value model, 0.0219 s^2^ for the constant firing rate model, and 0.0249 s^2^ for the trend model; SEM *<*0.0001 s^2^ in each case; mean fivefold cross-validated accuracy, 10 repeats, *N* = 37 neurons and 12 387 trials), indicating that the population-level activity patterns of serotonin neurons are best understood as a code for value, consistent with our previous work (12).

The aggregate performance statistics presented above conceal significant heterogeneity in the activity patterns of serotonin neurons. Some neurons exhibited trial-to-trial fluctuations in inter-trial firing that are clearly consistent with value (Fig. 3C), while other individual neurons had a seemingly constant firing rate, and still others exhibited a slow decrease in firing over the course of the recording session that visually resembles a linear trend (Fig. 3E). Under our value coding model, this heterogeneity is explained in terms of variability in 1) the strength of the effect of value on firing (termed normalized firing rate gain 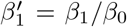) and 2) the learning timescale *τ*. The heterogeneity in learn-ing timescale spanned a large range, from 5 min to almost one hour, with the majority of neurons displaying a learning timescale between 17 and 33 min (100-200 trials). The non-overlapping credibility intervals in Fig. 3E shows that for those neurons, the learning timescale was different.

Across the different cells, the distribution of maximum a posteriori estimates (MAP) of the correlation with value coding (*β*_1_) shifted to higher gain from the distribution used as a prior (Fig. 3F). This indicates a general tendency of cells to correlate with value estimated at different timescales. More specifically, to calculate the fraction of cells that were positively correlated with value, we computed the MAP distribution for the fraction of cells having a gain *β*_0_ larger than zero. We found a distribution concentrated around 80% of the cells (Fig. 3G). These estimates are show more consistent value coding than could be extracted from a previous study using classical trace conditioning (9), as shown in gray in Fig. 3F-G.

Turning to the statistics of the learning timescale across cells, Figure 3H shows samples from the prior and the posterior for the learning timescale parameter. As illustrated in Fig. 3D, the posterior estimates concentrated close to a timescale corresponding to 100 trials. This is made more precise by computing the posterior distribution for the mean learning timescale of the full population of serotonin neurons. Our posterior peaked sharply at a value of 125 trials, such that the average estimation timescale across the serotonergic population matches quite closely the estimate of 130 trials of similar duration from Ref. (15).

## 3 THE ROLE OF THIRST AS A CONFOUNDING FACTOR

It is possible, however, that confounding factors are integrated in the inference of the timescale and associated value code potency. We thus consider a control variable that relates to elapsed time or thirst, which we refer to as ‘thirst’ for short. It is defined as an inverse tally of the number of water rewards consumed (Section 5.2.2). This type of of variable can influence motivation, behaviour, and perception of reward in this task. Although the rate of reward collection, and therefore the rate of thirst reduction, fluctuates over the course of the experiment, the overall time-course of thirst generally resembles a linear trend (Fig. 4B). As expected, this means that if the learning timescale *τ* is large, value and thirst closely resemble each other (Fig. 4C), leading to a high degree of correlation/confounding between the two variables when *τ* ≥45 trials (Fig. 4D). Thus, trend-like activity in a subset of serotonin neurons identified by our value coding model may actually be due to the effects of thirst. The timecourse of modeled thirst and value estimated on a long timescale are so similar, we sought to estimate what proportion of the neuron encode value in a manner that cannot be account for by thirst. To this end, we developed a second Bayesian model that can assess the degree with which a given neuron correlates with value or if its activity can be equally explained by value or thirst (see Methods; Mixture definition).

**Figure 4.**
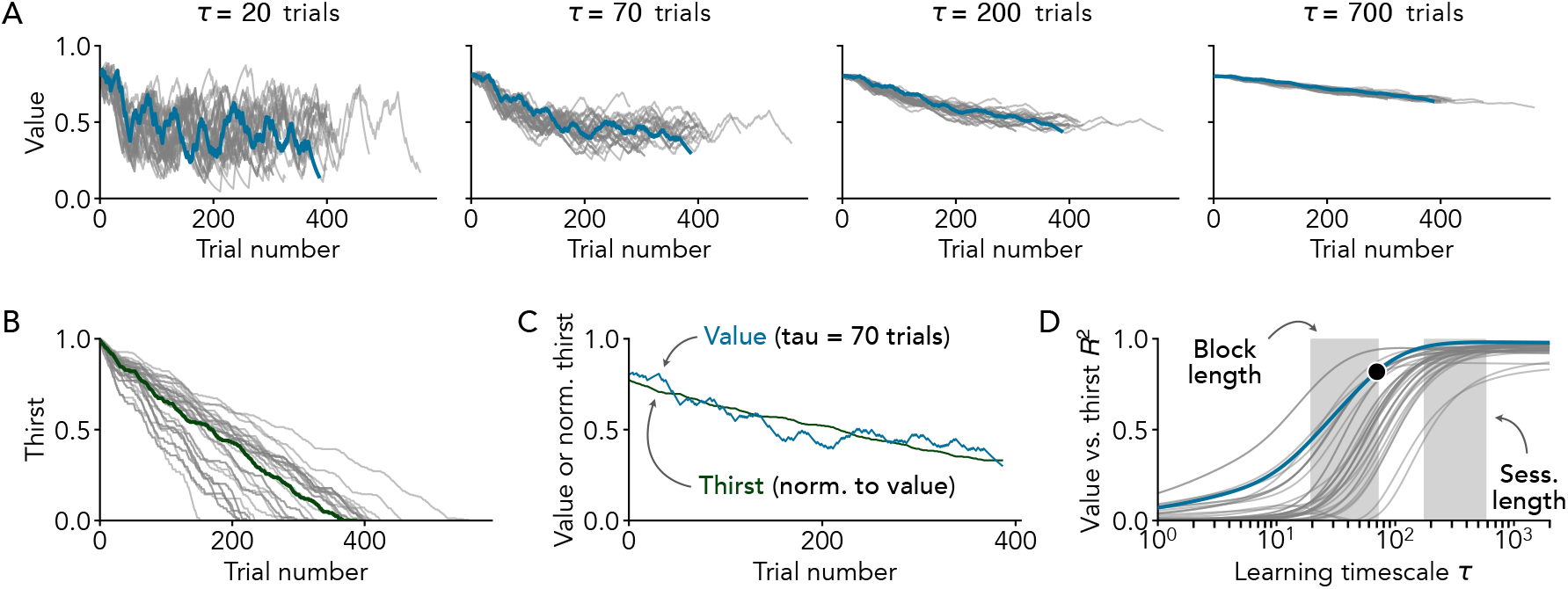
Value is confounded with thirst when learning is slow. **A** Dynamics of value as a function of the learning timescale τ. Each line represents one experimental session. An exemplar session is highlighted in blue. **B** Dynamics of thirst, defined as a quantity that decreases each time the animal consumes a water reward, starting at one and decreasing to zero over the course of a session. Each line represents one experimental session. The exemplar session from A is highlighted in green. **C** Comparison of value and thirst for the exemplar session highlighted in A and B. Value (blue) is shown using a learning timescale of τ = 70 trials. Thirst (green) is normalized so that its mean and standard deviation match those of value. **D** Confounding between value and thirst as a function of the learning timescale τ. Confounding is measured using the squared coefficient of determination, representing the fraction of variance in value that can be explained by thirst (and vice-versa). Each line represents one session. The exemplar session from A and B is highlighted in blue, and the black dot corresponds to the curves shown in C.

Fitting this mixture model to our data, we estimated that approximately half of neurons encode value (posterior mean 48.1 %, 95 % CrI 27.3–69.9 %), while a large minority encode thirst (posterior mean 41.7 %, 95 % CrI 21.0–63.1 %) and a small fraction are null coding (posterior mean 10.2 %, 95 % CrI 0.5–26.5 %; see Fig. S1B). The high level of uncertainty on each of these estimates is mainly due to the moderate number of neurons in this dataset rather than statistical confounding between value, thirst, and null coding (for comparison, a frequentist 95 % CI on a proportion of 50 % would be 33.7–66.3 % for *N* = 37 even in the absence of confounding). Null coding is less common than value and thirst coding (posterior probability 97.4 %, prior probability 33.3 %), but it is unclear whether value coding is more common than thirst coding or vice versa (posterior probability that value coding is more common 61.2 %, prior probability 50.0 %). We conclude that a large fraction of serotonin neurons encode value, but that a larger scale experiment is needed to determine whether this is the most common tuning feature.In the subset of serotonin neurons that unambiguously encode value, we estimated that rewards are typically integrated over several dozen trials in order to estimate value (posterior mean learning timescale 72.9 trials, 95 % CrI 45.1–132.5 trials; Fig. S1C). We still observed a very high level of variability in this feature across neurons, with the timescale over which serotonin neuron activity integrates rewards spanning roughly an order of magnitude (posterior mean standard deviation of the learning timescale 0.289 log10 units, 95 % CrI 0.136–0.558; Fig. S1D). Specifically, we estimate that the fastest 10 % of value coding serotonin neurons integrate rewards over fewer than 31.1 trials (posterior mean, 95 % CrI 14.2–52.3 trials; Fig. S1E) while the slowest 10 % integrate rewards over more than 170.8 trials (posterior mean, 95 % CrI 85.0–574.2 trials; Fig. S1F), with 80 % of neurons falling in the middle. Consistent with our model-free analysis (Fig. 2D and E), we estimate that 76.2 % (posterior mean, 95 % CrI 50.1–94.7 %) of value coding neurons integrate rewards over a period longer than the average block length in this experiment (45 trials). In sum, as one would expect by the act of removing those cells which are associated with a slow value estimation, the presence of thirst as a confounding factor reduces the proportion of neurons that unambiguously code for value and their estimation timescale is shorter.

### 3.1 Anticipatory licking depends on thirst and short-term reward history

We next sought to identify possible behavioural correlates of serotonin neuron activity. To determine whether longterm reward history or thirst influence the rate of anticipatory licking, we fitted a model in which licking is driven by a latent variable that reflects a mixture of value estimated over three different timescales, a fast timescale (1 trial), a medium timescale (10 trials) and a slow timescale of which matches the distribution of timescales observed in serotonin neurons (Fig. S1), and thirst (illustrated in Fig. 5A). Importantly, neither the fast nor the medium timescale match that observed in serotonergic cells (shortest timescale in Fig. 3D is 30 trials). Since this experiment contains both normal rewarded trials and a small number of cued unrewarded catch trials, we further modeled the lick rate as being dependent on the animal’s perception of trial type (see Methods for details). Animals correctly perceived the trial type 93.6 % of the time (posterior mean, 95 % CrI 88.3–96.8 %, population parameter estimated using a hierarchical model fitted to 9693 trials across 28 sessions), licking in response to the cue for rewarded trials but not in response to the cue for catch trials. The fact that animals correctly perceived catch trials and adjusted the rate of anticipatory licking accordingly confirms that this behaviour is at least partially related to value.

**Figure 5.**
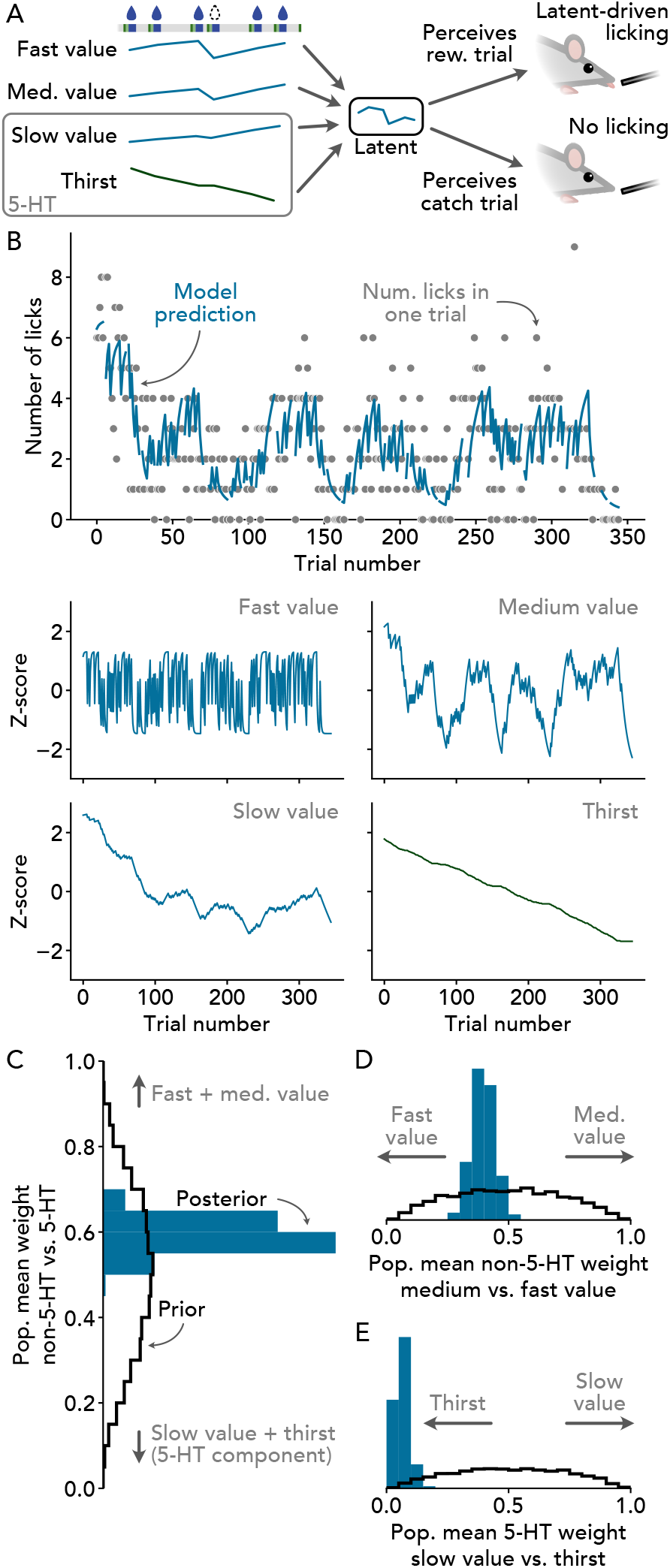
Analysis of anticipatory licking. **A** Overview of behavioural model. Value computed based on a short reward history (fast value, τ = 1 trial), medium reward history (medium value, τ = 10 trials), long reward history (slow value, τ chosen to match distribution found in serotonin neurons, mean of approximately 72 trials), and thirst are combined to form a latent that drives anticipatory licking if the animal believes it is in a r egular, rewarded trial rather than a catch trial. See Methods for details. **B** Data, MAP model prediction, and corresponding latent components for an exemplar session. For clarity, perceived catch trials are not shown (note gaps in blue line). **C** Mean weight assigned to 5-HT-related (slow value and thirst) vs. non-5-HT-related (fast and medium value) latent components across N = 28 sessions. **D** Mean weight assigned to medium value vs. fast value across N = 28 sessions. **E** Mean weight assigned to slow value vs. thirst across N = 28 sessions.

The fitted behavioural model successfully extracted the main features of anticipatory licking visible in the raw data (Fig. 5B). In particular, fitted model predictions exhibited large modulations that reflect the block structure of the experiment, session-wide decreasing trends that may reflect satiety/thirst, and trial-to-trial fluctuations that reflect very short-term reward history. The first two of these features are consistent with our model-free analysis that pointed to anticipatory licking evolving over two timescales, one that reflects block structure and another that reflects a long-term trend (Fig. 2). Overall, our model explained the majority of the observed variation in the rate of anticipatory licking (67.2 ± 14.3 % variance explained by MAP parameter estimates, mean ± SD, *N* = 28 sessions), further confirming that this behaviour depends on some mixture of value and thirst, while leaving open the possibility that other factors may play an important role.

Using the parameters estimated by our behavioural model, we next sought to determine the relative contributions of the value and thirst factors to anticipatory licking. We found that factors consistent with serotonin neuron activity, namely slow value and thirst, were collectively given a weight of 40.8 % (95 % CrI 34.0–47.7 %) even though these factors both evolve over timescales considerably longer than the block length in this experiment. Within the nonserotonergic factors (collectively weighted at 59.2 %, 95 % CrI 52.3–66.0 %; see Fig. 5C), fast value was given slightly more than half of the weight (60.4 %, 95 % CrI 51.1–69.4 %; see Fig. 5D), indicating that anticipatory licking depends meaningfully on value estimated over short and medium timescales. Within the serotonergic factors, the overwhelming majority of the weight was given to thirst (93.6 %, 95 % CrI 87.2–97.7 %; see Fig. 5E), indicating that value estimated over a timescale significantly longer than the average block length correlates less with anticipatory licking than our simple model of thirst. Taken together, anticipatory licking is explained partially by value integration occurring at a faster time scale than seen in serotonergic cells and partially by value integration happening at a timescale that matches the serotonergic cells, but the part of the behaviour matching the timescale seen in sertonergic cells is better explained by a model of thirst than a model of value.

## 4 DISCUSSION

We have reported a Bayesian model fitting approach for how individual serotonin neurons encode the evolving state value over time. Our model accounts for additional factors such as thirst or elapsed time and reveals that neurons vary in how strongly their firing reflects state value and in the timescales over which they update this value, with an average learning timescale of between 10 and 20 minutes. By also analyzing simultaneous behavioral data, specifically anticipatory licking, we find that behaviour is in most part linked to value signals learned on faster timescales than those encoded by serotonin neurons, while the part that is consistent in the serotonergic estimation timescale can be entirely explained by thirst as a confounding factor. These results offer a detailed quantification of how individual serotonin neurons represent value and its learning dynamics, uncovering a mismatch between the value estimation timescales in serotonergic cells and those influencing behaviour.

### Relationship between neural activity and behaviour

There is a significant amount of variation in both neural activity and behaviour that is not explained by the models considered here. We have not attempted to determine whether the unexplained variation is correlated between neural activity and behaviour in the corresponding sessions, leaving open the possibility that some signal other than thirst or value may be responsible for driving both neural activity and behaviour. In principle, we could figure this out by looking for correlations in the residuals of the neural activity and behavioural models, but we determined that this was not feasible due to technical challenges. In particular, the fact that the residuals in the neural model depend on the coding scheme used by each neuron, which is treated as uncertain in our approach, and the fact that the experiment contains catch trials which give the behavioral residuals a structure that is very difficult to model.

### Value of the trace period period

We have compared trial-to-trial dynamics of anticipatory licking, which mostly takes place in the trace period, with the trial-to-trial fluctuations of serotonergic neuron activity, which were recorded during the inter-trial period. This approach is supported by the prospective value theory, where the value in the inter-trial period scales with the value in the trace period. This is statements, however, arises from the dynamics of value after learning. The dynamics of value during learning depends heavily on the type of learning algorithm that is being used. For example, the use eligibility traces in the learning algorithm will alter the speed of value learning. Importantly, some learning algorithms can, in principle, make the learning timescale of periods far from the reward different from that close to the reward. Experimental observations of Ref. (12) do not support this possibility as, on a 5 trial horizon, the speed at which the trace period activity of serotonin neurons changed matched the speed measured in the inter-trial period.

### Estimation timescale of dopaminergic activity

Matias et al. (15) found that the reward associations reflected in bulk serotonergic activity changed over the course of approximately 130 trials. This was an order of magnitude slower that the reward associations reflected in bulk dopaminergic activity, which changed over the course of approximately 20 trials (see also Ref. (10)). Given that their trials were about the same duration as the trials in the experiment analyzed here (9-14 s in Ref. (15) vs. an average of 9 s in Ref. (16)), we can expect that dopaminergic activity would change over the time course of 20 trials in the mice recorded here. It is interesting to note that this learning timescale, by beeing close to the medium timescale used in Fig. 5, showed some correlation with anticipatory (Fig. 5D), suggesting that the value estimation timescale reflected in dopaminergic cells is more closely associated with behaviour than the value estimation timescale seen in serotonergic cells (15).

### Capturing activity fluctuations in vivo

Mathematical models have been developed for predicting activity fluctuations of single neurons in vitro (18–24) and in vivo (25–28). While it is difficult to compare model performance across different recording modalities and different metrics (29), we note that models of cortical neurons tend to achieve high prediction accuracy in slice (20, 21, 30, 31) but very low prediction accuracy in vivo (25). This discrepancy may be attributed to many missing latent factors (27, 32), population multiplexing of information (33–35) as well as the difficulty of capturing the relationship with cortical cell’s inputs (36–42). Serotonergic neurons on the other hand are not particularly well captured by integrate-and-fire models in vitro (24). In this context, the variance explained achieved by our model for predicting activity fluctuations in vivo is surprisingly good. In contrast to the cortex, serotonergic neurons of the dorsal raphe are not numerous and may the population may not be able to support population multiplexing (33). These neurons are also much less bursty and cortical neurons (43), indicating that they may also not be able to support burst multiplexing (35).

## 5 METHODS

### 5.1 *In vivo* experiment

The Pavlovian conditioning experiment with dynamic reward probabilities has been reported previously by Grossman et al. (16). A brief summary follows.

Odour-cued water rewards were delivered to five water-restricted C57/BL6 mice (four male, one female) over the course of 28 recording sessions, each of which lasted ~ 1 h.Rewards were delivered stochastically with a probability of 0.2, 0.5, or 0.8 which changed dynamically according to a block structure. Blocks lasted 20–70 trials (uniform distribution), and the first block of each session always had a reward probability of 0.8. Catch trials consisting of a different odour cue followed by no reward were randomly interleaved with regular trials throughout the experiment at an average rate of 1*/*20 trials. Catch trials and trials in which a reward was available but not collected were deemed unrewarded for the purposes of our analysis. Mice collected 0.45 ± 0.04 rewards per trial and completed 356 ± 88 trials per session (mean ± SD across sessions). Inter-trial intervals followed an exponential distribution with mean 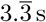. Spike times of optogenetically-tagged serotonin transporter-expressing neurons were collected using tetrode recordings at a rate of 1–3 neurons per session (mean 1.32 neurons).

Experimental and surgical procedures were performed in compliance with the *National Institutes of Health Guide for the Care and Use of Laboratory Animals* and were approved by the Animal Care and Use Committee at Johns Hopkins University.

### 5.2 Models

#### 5.2.1 Value

We define value as the expected reward in the upcoming trial, estimated using the Rescorla–Wagner learning rule

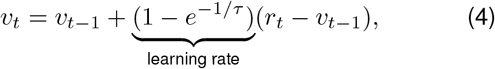

where *v*_*t*_ is the value during the inter-trial interval immediately following the reward *r*_*t*_ on trial *t*, and *τ* is the learning timescale.

The model is written in terms of the learning timescale *τ* instead of the more conventional learning rate *α* because the learning timescale can be interpreted as the number of trials over which past rewards are integrated in order to estimate the expected reward in the upcoming trial. However, it is important to note that the two are equivalent: τ = −1/ ln(1 − α)

#### 5.2.2 Thirst

We define thirst as a quantity *h*_*t*_ that decreases by a fixed amount each time the animal consumes a water reward.

We arbitrarily set *h*_0_ = 1 and *h*_*T*_ = 0, where *T* is the number of trials, such that

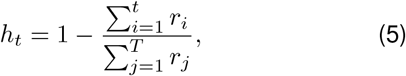

where *r*_*x*_ ∈{0, 1} is the binary water reward received on trial *x*. Reward collection had the tendency to follow the time of since the start of the experiment, but showed individual differences across mice Fig. S3.

#### 5.2.3 Latent variable model of neural activity

We model neural activity 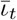 in terms of a latent firing rate *ρ*_*t*_ driven by a latent input *x*_*t*_. We assume a linear effect of the input on the firing rate

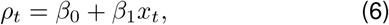

where *β*_0_ is the baseline firing rate when the input is zero and *β*_1_ is the gain. In turn, we model the mean inter-spike interval 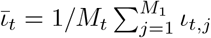, where *ι*_*t*,1_, *ι*_*t*,2_, …, *ι*_*t,M*_ are individual inter-spike intervals in trial *t*, in terms of the inverse firing rate, or frequency 1*/ρ*_*t*_

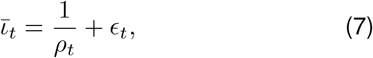

where *ϵ*_*t*_ is the residual at time *t* due to a combination of spiking noise, measurement noise, and drift. This quantity could only be estimated when there were at least two spikes in the data (Fig. S4). We found that this method was more stable in such sparse firing than using spike trains and their associated likelihood functions (as in Refs. (20, 21, 24, 44)).

#### Residual models

To simplify comparisons with null models based on constant firing rates or linear time trends (see Section 2.2), for the purposes of cross-validation we model the residuals *ϵ*_*t*_ as Gaussian white noise with time-varying variance

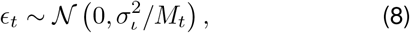

where *σ*_*ι*_ is the standard deviation of a single inter-spike interval and *M*_*t*_ is the number of inter-spike intervals in trial *t* (such that 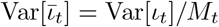).

During inference, we model the residuals according to a third-order autoregressive process in order to properly account for drift (Fig. S5). The process is defined as

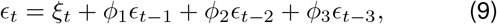

where 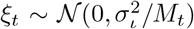 is Gaussian white noise driving the residuals and *ϕ*_*j*_ are weights constrained to [−1, 1].

In both cases, we use Gaussian noise as the basis of our residual model. We have chosen Gaussian noise for two reasons: 1) inter-spike intervals recorded *in vitro* in identified serotonin neurons are approximately Gaussian (data not shown), and 2) the Lyapunov central limit theorem guarantees that the *mean* inter-spike interval 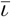 that is the focus of our analysis converges to a Gaussian distribution (with variance proportional to 1*/M*_*t*_) under mild conditions.

### 5.3 Data analysis

#### 5.3.1 Quantification of anticipatory licking

We measure anticipatory licking as the number of licks during the 2 s period between cue and reward onset.

#### 5.3.2 Quantification of neural activity

Neural activity is measured as the mean inter-spike interval 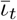 during each inter-trial interval. For inter-trial intervals with fewer than two spikes, the mean inter-spike interval is considered missing data.

#### 5.3.3 Estimation of power spectra

To estimate the power spectra of reward collection, anticipatory licking behaviour, and serotonin neuron firing activity, we consider each of these as stochastic processes and extract the power spectra as the smoothed Fourier cosine transforms of the corresponding mean sample autocorrelation functions (45), ignoring missing data.

For example, taking *x*_1,[*i*]_, *x*_2,[*i*]_, …, *x*_*T*_[*i*]_,[*i*]_ to be the number of anticipatory licks on trials 1 to *T*_[*i*]_ for mouse *i*, we compute the autocorrelation at lag *k* up to 400 as

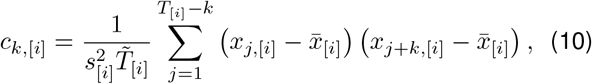

where 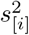 is the sample variance of 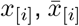 is the sample mean, and 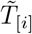 is the number of (*x*_*j*,[*i*]_, *x*_*j*+*k*,[*i*]_) pairs for which neither *x*_*j*,[*i*]_ nor *x*_*j*+*k*,[*i*]_ are missing values. Note that we take the term inside the sum to be zero if either *x*_*j*,[*i*]_ or *x*_*j*+*k*,[*i*]_ is missing, and we consider *c*_*k*,[*i*]_ itself to be missing if 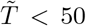. In the case of anticipatory licking, we treat the number of licks during catch trials as missing data. For firing activity, we treat the mean inter-spike interval during inter-trial intervals with fewer than two spikes as missing data. Reward data is never missing.

Given the sample autocorrelations of anticipatory licking *c*_*k*,[1]_, *c*_*k*,[2]_, …, *c*_*k*,[*N*]_ for *N* = 28 recording sessions, we estimate the underlying autocorrelation function of the licking process as the mean of the sample autocorrelations

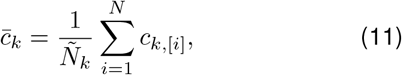

where 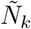 is the number of *c*_*k*,[*i*]_ that are not missing, and where the term inside the sum is taken to be zero if *c*_*k*,[*i*]_ is missing, as above. For firing activity, *N* = 37 neurons rather than 28 sessions.

Finally, the power spectrum is estimated from the autocorrelation function 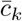 using the smoothed Fourier cosine transform

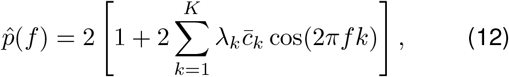

where 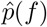 denotes the estimated power at frequency *f* rather than a probability (following the notation of ref. 45), *K* = 399 is the maximum lag, and the *λ*_*k*_ are smoothing coefficients. The smoothing coefficients are set according to *λ*_*k*_ = exp[−(*k*^4^*/*(*K/*2.5)^4^)], which is close to one for lags less than 100 and close to zero for lags greater than 200.

#### 5.3.4 Hierarchical Bayesian model of value coding Notation

This section describes a model of data from multiple neurons. Sub-scripts in square brackets _[·]_ are used to indicate quantities specific to a particular neuron. For example, *τ*_[*i*]_ is the learning timescale for neuron *i*.

##### Posterior definition

The log posterior density *𝓁*_*p*_ of our model of serotonin neuron activity is defined as

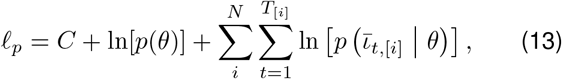

where *C* is a constant term that represents the log probability density of the data (the denominator in Bayes’ rule), *p*(*θ*) is the prior, *N* = 37 neurons, *T*_[*i*]_ is the number of trials for neuron *i*, and 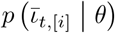 is the likelihood of the mean interspike interval for neuron *i* on trial *t* given the parameters *θ*. Whenever 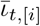 is missing, we take the corresponding likelihood to be one.

##### Parameterization and priors

The choice of model parameterization has a dramatic effect on whether/how quickly the Hamiltonian Monte-Carlo algorithm used for Bayesian inference will converge (46), but does not affect the inferences that can be drawn about quantities of interest at convergence. For technical reasons, we have parameterized the model in terms of the following quantities:

- The learning timescale on a log10 scale log *τ*_[*i*]_.
- Normalized initial value for each neuron 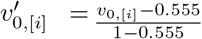.
- Firing rate intercept population log shape and scale parameters ln *a*, ln 1*/b*.
- Firing rate intercept for each neuron *β*_0,[*i*]_.
- Normalized firing rate slope for each neuron 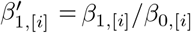.
- Estimated ISI standard deviation for each neuron *σ*_*τ*,[*i*]_.
- Drifting noise process coefficients for each neuron *ϕ*_1,[*i*]_, *ϕ*_2,[*i*]_, *ϕ*_3,[*i*]_ (only for inference).

Priors for all parameters are listed in Table 1.

**Table 1.**
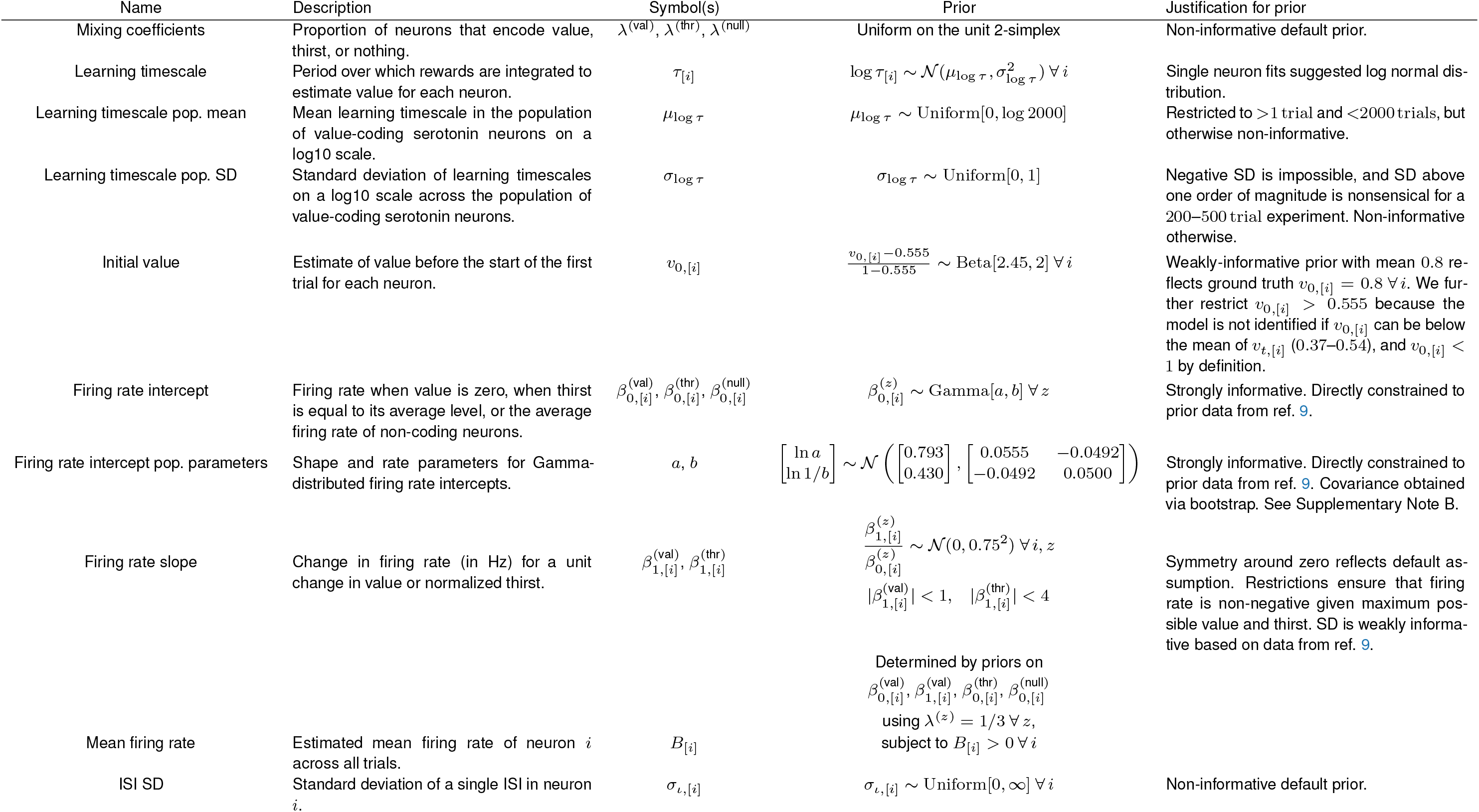
Summary of quantities of interest in the hierarchical Bayesian mixture model of serotonin neuron activity.

##### Likelihood

The likelihood 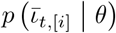 is derived by substituting the value *v*_*t*_ from Eq. (4) into the inter-spike interval model given by Eq. (7) evaluating the probability density of the residuals *ϵ*_*t*_ under the appropriate noise model. For cross-validation, we use the Gaussian white noise given in Eq. (8). For Bayesian inference, we use the drifting noise process given in Eq. (9).

#### 5.3.5 Hierarchical Bayesian mixture model of neural activity

##### Notation

This section describes a model of data from multiple neurons in terms of multiple possible coding schemes. In addition to sub-scripts in square brackets used to indicate quantities specific to a particular neuron, superscripts in round brackets ^(·)^ are used to indicate quantities or functions specific to a particular coding scheme. For example, *p*^(val)^ (*x*_[2]_) denotes the probability density of *x* from neuron 2 under the value coding scheme.

##### Mixture definition

To measure the value and thirst coding features of serotonin neurons, we fitted a mixture model in which the time-dependent activity of each neuron 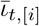 reflects either value *v*_*t*,[*i*]_, thirst *h*_*t*,[*i*]_, or nothing (i.e., the underlying firing rate is constant), with corresponding likeli-hoods 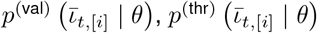, and 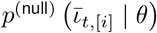. Given the prior *p*(*θ*), the overall log posterior density of the mixture model is therefore given by

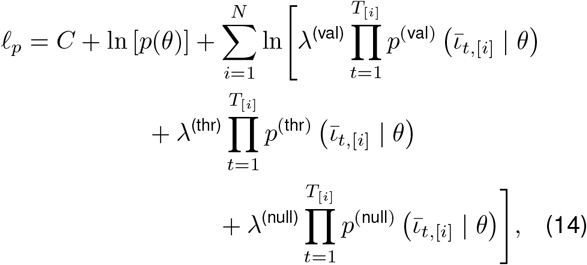

where *C* is a constant that represents the log probability density of the data (the denominator in Bayes’ rule), *N* = 37 neurons, *T*_[*i*]_ is the number of trials for neuron *i*, and the *λ* are mixing coefficients on the unit simplex Σ_*z*_ *λ*^(*z*)^ = 1. When 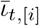 is missing, we take the corresponding likelihoods to be 1. Details on the prior and likelihoods are given below.

##### Parameterization

The mixture model is parameterized in terms of the following quantities:

- Mixing coefficients *λ*^(val)^, *λ*^(thr)^, *λ*^(null)^.
- Learning timescale population parameters *µ*_log *τ*_, *σ*_log *τ*_.
- Standardized learning timescale for each neuron 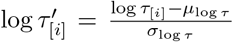. This parameterization is chosen to avoid a Neal’s funnel effect when *σ*_log *τ*_ is small (46).
- Normalized initial value for each cell 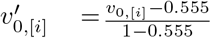.
- Firing rate intercept population shape and scale parameters ln *a*, ln 1*/b*.
- Estimated mean firing rate for each neuron *B*_[*i*]_.
- Normalized firing rate slopes for each neuron under the value and thirst models 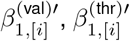, where 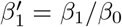
- Estimated ISI standard deviation for each neuron *σ*_*ι*,[*i*]_.
- Drifting noise process coefficients for each neuron *ϕ*_1,[*i*]_, *ϕ*_2,[*i*]_, *ϕ*_3,[*i*]_ (only for inference).

##### Firing rate intercepts

The firing rate intercepts 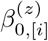 are an important source of prior information in our model (see Table 1) even though they do not appear directly in its parameterization (see above). Under the value model, the firing rate intercept is derived from the estimated mean firing rate *B*_[*i*]_ and normalized firing rate slope 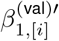 according to

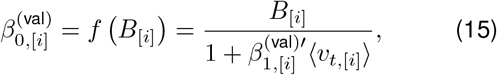

where *v*_*t*,[*i*]_ is the mean value across trials with at least one ISI for neuron *i*. Note that the absolute derivative of this transformation is

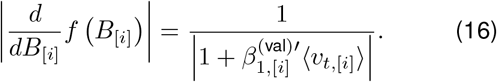

Under the thirst and null models, the firing rate intercept is equal to the baseline firing rate

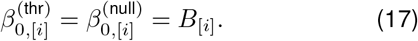

To incorporate prior information about the firing rate intercepts 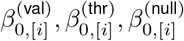 into our model, we need to decompose the overall prior *p*(*θ*) into a set of terms that depend on the *β*_0_. First, we factorize *p*(*θ*) to obtain the prior probability of the mean firing rates *B*_[*i*]_ conditional on the remaining parameters *θ*^c^

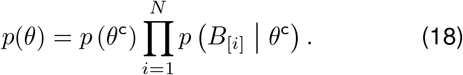

Next, we marginalize the conditional probability of the mean firing rate over the coding schemes

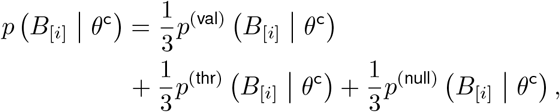

where we have assigned each of the three coding schemes an equal prior probability of 1*/*3. Using Eqs. (15) to (17), the above can be rewritten in terms of the *β*_0_ under each model

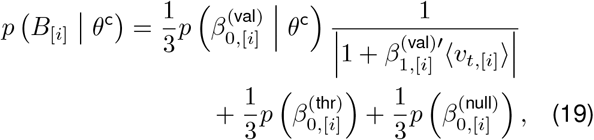

where the first term includes an adjustment of 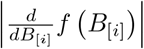 to account for the change of variables from *B*_[*i*]_ to 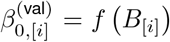. The second and third terms do not need an adjustment for the change of variables, nor do the slopes depend on other parameters *θ*^c^.

##### Priors Using Eqs. (18) and (19), the overall prior for the mixture model can be written as

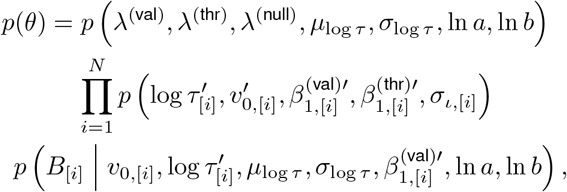

where all of the parameters are conditionally independent except for *λ*^(val)^, *λ*^(thr)^, *λ*^(null)^ due to the simplex constraint. The exact form of the prior on each parameter is listed in Table 1.

##### Value likelihood

The likelihood under the value model 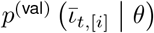 is derived by substituting *v*_*t*,[*i*]_ from Eq. (4) (using *v*_0,[*i*]_ for the initial condition) in for *x*_*t*_ in Eq. (6), yielding

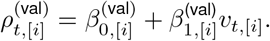

The firing rate 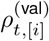 is then substituted into the inter-spike interval model given in Eq. (7), yielding

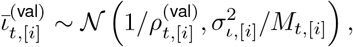

which defines the Gaussian likelihood of 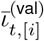.

##### Thirst likelihood

The likelihood under the thirst model 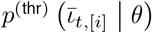 is derived analogously to the value likelihood described above, using *x*_*t*_ = (*h*_*t*,[*i*]_ − ⟨*h*_*t*,[*i*_] ⟩)0.381 as the input to the firing rate model from Eq. (6). The thirst signal is centered so that 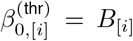 and scaled by a factor of 0.381 to approximately match the average dynamic range of value *v*_*t*,[*i*]_ using *τ*_[*i*]_ = 70 trials and *v*_0,[*i*]_ = 0.8 ∀ *i*.

##### Non-coding likelihood

The likelihood under the noncoding model 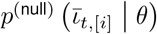 is derived analogously to the value and thirst likelihoods using *x*_*t*_ = 0 ∀ *t* in Eq. (6).

#### 5.3. Hierarchical Bayesian model of anticipatory licking

##### Notation

Similar to previous sections, we use subscripts in square brackets ·_[*i*]_ to denote quantities that are specific to session *i*.

Since this experiment contains a mixture of regular trials with stochastic rewards and catch trials in which reward was delivered, we model the number of anticipatory licks in trial *t* of session *i, l*_*t*,[*i*]_, as being dependent on whether the animal perceives that it is in a regular or catch trial. To indicate this dependence, we use the indicator *z*_*t*,[*i*]_ = 1 to denote the event that the animal is in a regular trial and 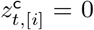 to denote the event that the animal is in a catch trial. Similarly, we use 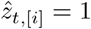 to indicate that the animal *perceives* that it is in a regular trial, and 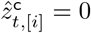 to indicate that the animal *perceives* that it is in a catch trial.

### Perceptual model of licking

We model the number of anticipatory licks *l*_*t*,[*i*]_ a nonlinear function of a standard-ized latent variable 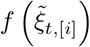 (defined below) if the animal perceives that it is in a regular trial, and as being near zero if the animal perceives that is in a catch trial

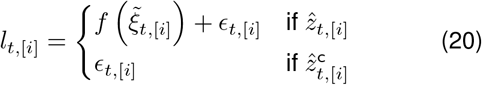

where 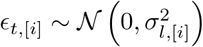 is noise. The log posterior probability density of the licking model is

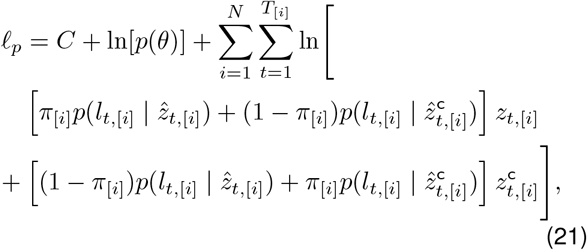

where *C* is a constant that represents the log probability of the data; *p*(*θ*) is the prior; *N* is the number of recording sessions; *T*_[*i*]_ is the number of trials in session *i*; *π*_[*i*]_ is the probability that the animal perceives the trial type correctly in session *i*; and the likelihoods come from the normal densities implied by the noise terms in Eq. (20). We model the variability in *π*_[*i*]_ across sessions using a beta distribution

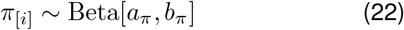

from which we can estimate the population mean correct perception probability reported in the main text.

### Licking in perceived regular trials

For perceived regular trials, we model the number of anticipatory licks as a non-linear function of a standardized latent variable 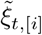 (defined below)

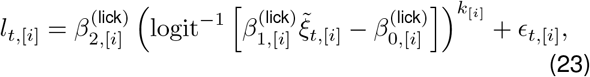

where logit^*⨪*1^[·] is the logistic function; 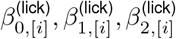 are the intercept, input gain, and output gain, respectively; and *k*_[*i*]_ is a parameter to control the shape of the nonlinear function (*k*_[*i*]_ *>* 0). The subscript in square brackets _[*i*]_ denotes a quantity that is specific to recording session *i*, analogous to the subscripts that denote neuron-specific quantities in the sections above.

For ease of fitting, we reparameterize the intercept 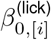 and input gain 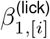 in the above model in terms of two anchor points *A*_1,[*i*]_, *A*_2,[*i*]_ that we constrain to be somewhere in the non-saturated range of the logistic function

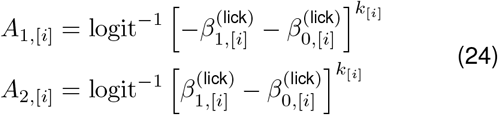

and use these to back-calculate the input gain 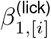 and intercept 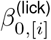. This procedure ensures that when the standardized latent 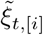 varies between *⨪*1 and 1, the output of the logistic function will vary between *A*_1,[*i*]_ and *A*_2,[*i*]_, which we constrain to 0 *< A*_1,[*i*]_ *<* 1*/*2 and 1*/*2 *< A*_2,[*i*]_ *<* 1.

To obtain the input gain and intercept from the anchor points, we first solve Eq. (24) for 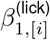, yielding

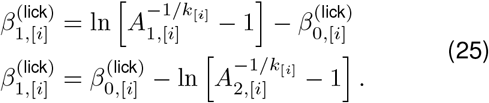

Subtracting these two equations and rearranging yields an equation for 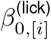

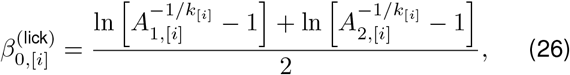

which can be substituted back into Eq. (25) to obtain 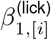.

#### Latent

The unstandardized latent variable *ξ*_*t*,[*i*]_ is defined as a mixture of thirst and value computed with three different learning timescales

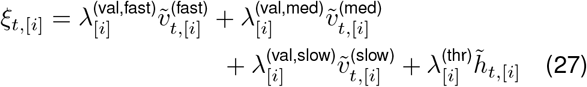

where 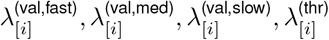 are mixing coeffi-cients constrained to the unit simplex (*i*.*e*., sum to one). We use a tilde 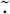 to indicate a standardized variable

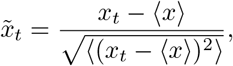

Where⟨ ·⟩ denotes the mean across trials, as above. Note that the variable 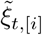 used to drive licking in Eq. (23) is a standardized version of Eq. (27). We model variability in the mixing coefficients across sessions using a Dirichlet distribution

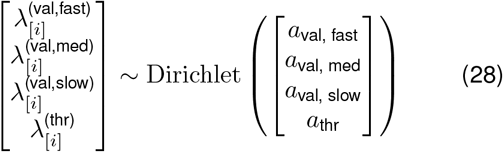

from which we can estimate the population mean of each mixing coefficient reported in the main text.

The unstandardized values *v*^(*x*)^ used to drive the latent are parameterized by the initial value 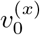 and learning timescale *τ* ^(*x*)^ (recall Eq. (4)). The initial value is taken from a beta distribution and scaled to lie between 0.555 and 1

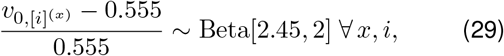

as in the neural activity models (Table 1). The fast values *v*^(fast)^ use *τ* = 1 trial and the medium values *v*^(med)^ use *τ* = 10 trials. The slow values *v*^(slow)^, which are intended to reflect the potential influence of value-coding serotonin neurons on behaviour, use learning timescales taken from a log-normal distribution that closely approximates the posterior population distribution of learning timescales from the neural data

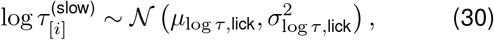

subject to 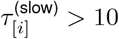, where

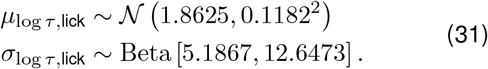

The parameters of the *µ* _log *τ*,lick_and *σ* _log *τ*,lick_ distributions_1,[*i*]_were obtained via maximum likelihood fits to the corre-_[*i*]_ sponding posterior distributions from the neural activity mixture model. This constrains the 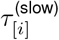 in the behavioural model to match the neural activity model, while accounting for uncertainty inherent in the neural model.

#### Parameterization and priors

For the purposes of fitting, the model is parameterized in terms of the following quantities:

- The correct perception probabilities *π*_[*i*]_ and their pop-ulation distribution parameters *a*_*π*_, *b*_*π*_.
- Normalized initial value for each session 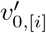, as in the neural activity models.
- Standardized slow learning timescale for each session 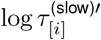 and corresponding population distribution parameters *µ*_log *τ*,lick_, *σ*_log *τ*,lick_, as in the hierarchical neural activity model.
- Mixing coefficients for each session 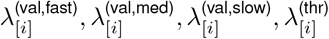 and corresponding population distribution parameters*a*_val, fast_, *a*_val, med_, *a*_val, slow_, *a*_thr_.
- Nonlinear function anchor points for each session*A*_1,[*i*]_, *A*_2,[*i*]_.
- Nonlinear function shape parameter on a log10 scale log *k*_[*i*]_ for each session.
- Nonlinear function output gain 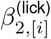 for each session.
- Lick count standard deviation for each session *σ*_*l*,[*i*]_.

---

**Figure S1.**
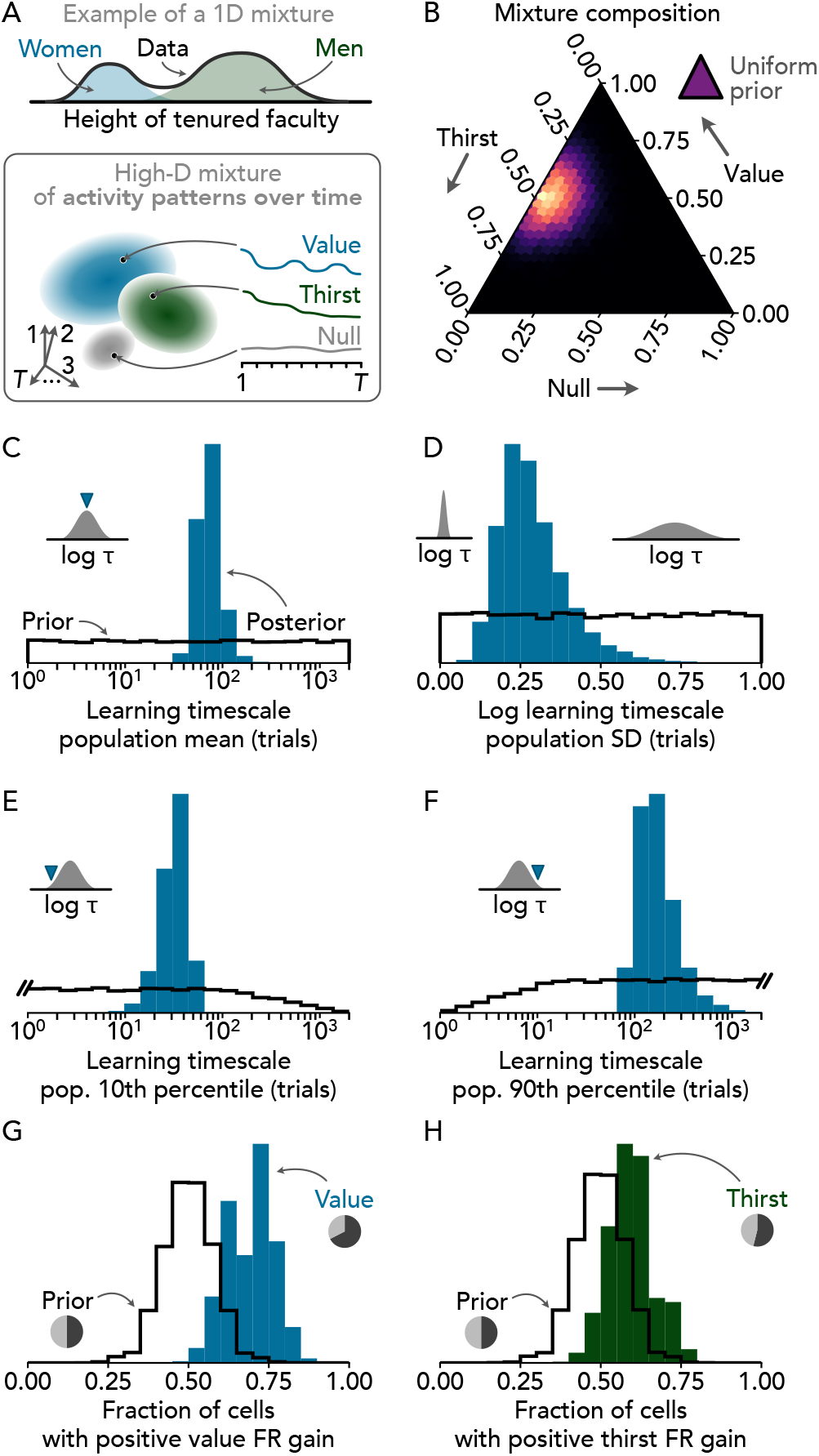
Features of value and thirst coding in serotonin neurons. **A** Conceptual introduction to the value, thirst, and null coding mixture model. A hypothetical one-dimensional dataset of hairline measurements in tenured faculty is illustrated at top. A mixture model can infer the average hairlines of male and female faculty as well as the proportion of faculty that are male or female, even when gender information is missing from the data. Similarly, the mixture model of serotonin neuron coding features can infer the properties of value, thirst, and null coding neurons as well as the proportion of neurons of each type (bottom). **B** Inferred proportion of neurons that are value, thirst, or null coding. Brighter colours in the main plot indicate higher posterior density. The uniform prior is illustrated schematically at top right. **C** Mean learning timescale *τ* across the population of value-coding serotonin neurons. **D** Standard deviation of the learning timescale *τ* across the population of value-coding serotonin neurons on a log_10_ scale. **E** 10^th^ percentile of the distribution of the learning timescale *τ* across the population of value-coding serotonin neurons. **F** 90^th^ percentile of the distribution of the learning timescale *τ* across the population of value-coding serotonin neurons. **G** Fraction of cells in this dataset with positive firing rate gain for value *β*^(val)^. Pie plot shows the MAP estimate of the proportion of neurons with positive (negative) firing rate gain in black (gray). Note that the prior (black line) is based on a default assumption that positive and negative gains are equally common in the population. **H** Fraction of cells in this dataset with positive firing rate gain for thirst *β*^(thr)^. Presented as in G. All parameter estimates are based on a model fitted to 12 387 trials from 37 neurons. Black lines and coloured histograms in C–H indicate the prior and posterior, respectively, for each parameter or derived feature. B–D show parameters that are directly included in the model, while E–H are derived from fitted parameters. Partial-pooling across neurons was used for the quantities shown in C–F, full pooling was used for the parameters in B, and no pooling was used for G or H. See Section 5.3.5 for details.

**Figure S2.**
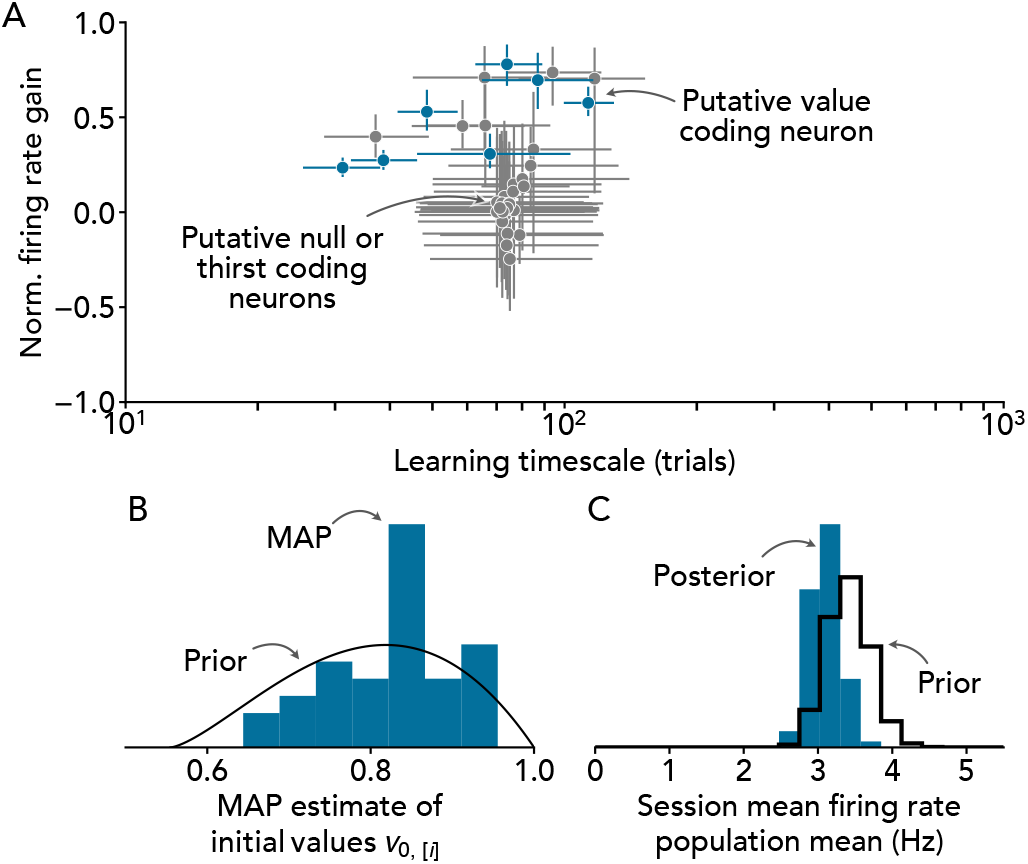
Additional features of value, thirst, and null coding mixture model. Related to Fig. S1. **A** Estimated normalized firing rate gain and learning timescale for each neuron. Points and error bars represent the median and IQR of the posterior. Neurons for which the 95 % credibility interval for the normalized firing rate gain includes zero are indicated in gray. **B** Distribution of estimated initial values *v*_0,[*i*]_ from the *maximum a posteriori* (MAP) sample (blue histogram) compared with the prior (black line). **C** Posterior distribution of mean firing rate in the population (blue) compared with the prior (black).

**Figure S3.**
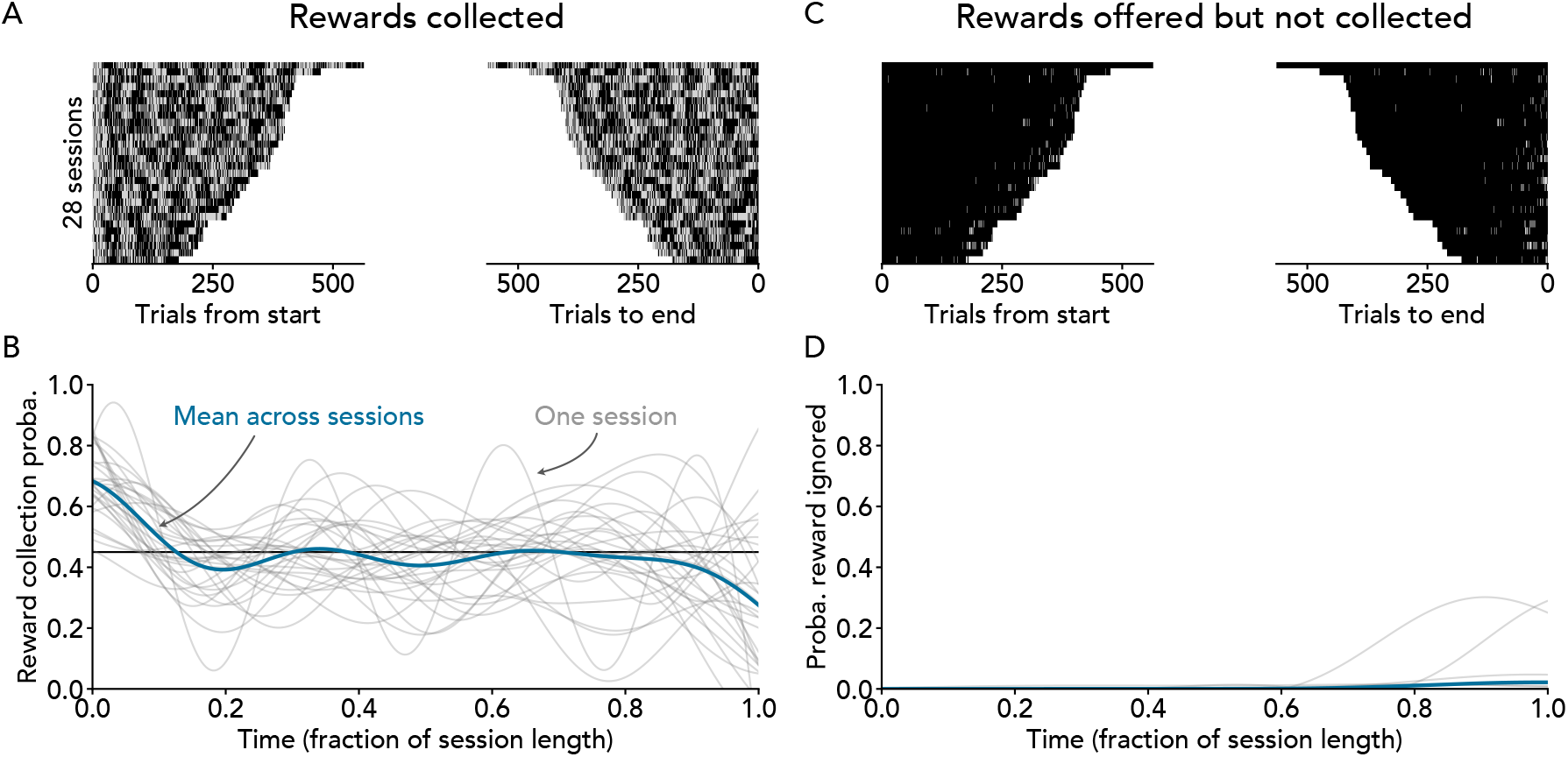
Time-course of reward collection across recording sessions. **A** Heatmaps showing whether or not a reward was collected on each trial, aligned to either the start (left) or end (right) of the experiment. Trials in which a reward was collected are shown in gray, no-reward trials are shown in black. **B** Time-course of reward collection aligned to fraction of session length. Each gray line represents data from one row of A smoothed using Gaussian process regression (observation variance set to 0.45 *×* (1 *⨪* 0.45), squared exponential kernel with length scale 1*/*6). Black line represents the expected mean reward probability of 45 %. Note that the reward collection probability is high at the start of the experiment because the first block of trials always has a reward probability of 80 %. **C** Heatmaps showing trials where a reward was offered but not collected (ignored rewards), aligned as in A. Trials in which a reward was offered but not collected are shown in gray, trials in which a reward was either not offered or offered and collected are shown in black. **D** Time-course of the rate of ignored rewards, presented as in B.

**Figure S4.**
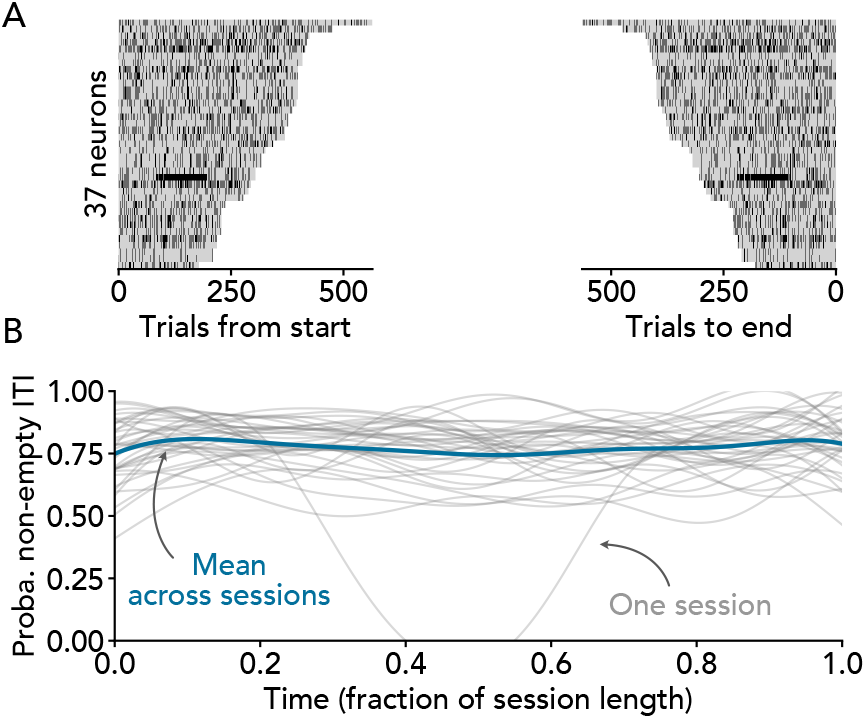
Time-course of inter-trial intervals (ITIs) with missing inter-spike interval (ISI) data across neurons. Missing ISI data is the result of ITIs with fewer than two spikes. This can occur if the ITI is shorter than the mean ISI (the minimum ITI duration is 0 s while serotonin neuron ISIs are typically approx. 0.2–1 s), or if the neuron is lost due to spike sorting or other recording issues. **A** Heatmaps showing whether or not each ITI has at least one ISI, aligned to either the start (left) or end (right) of the experiment. ITIs with zero ISIs are shown in black, ITIs with at least one ISI are shown in gray. **B** Time-course of missing ISI data aligned to fraction of session length. Each gray line represents data from one row of A smoothed using Gaussian process regression (observation variance set to *p ×* (1 *⨪ p*), where *p* is the fraction of non-empty it is in the corresponding session; squared exponential kernel with length scale 1*/*6).

**Figure S5.**
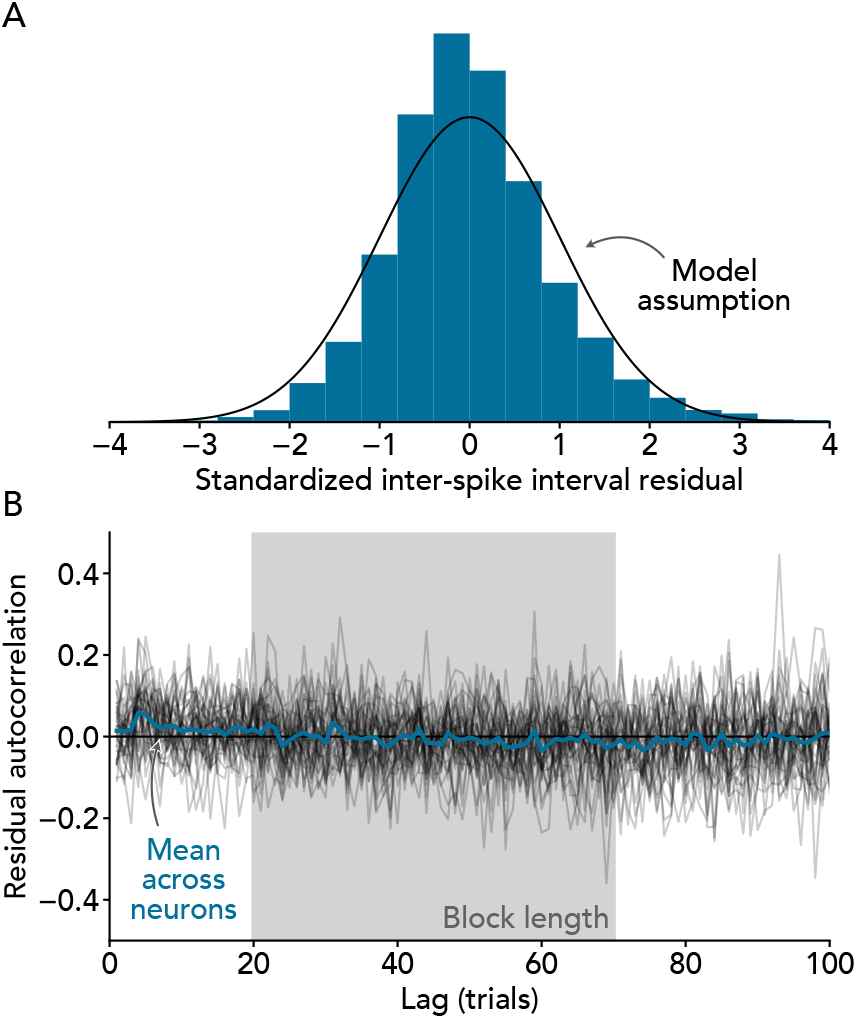
Value coding model residuals compared with model assumptions. **A** Histogram of standardized inter-spike interval residuals (blue) compared with the normal distribution assumed by the model (black line). **B** Autocorrelation of standardized residuals for each neuron (gray lines) compared with the model assumption of zero autocorrelation (black line). For clarity, the autocorrelation at zero lag is not shown. Residuals are net of drift captured using a third-order autoregressive model.

## A JUSTIFICATION FOR BAYESIAN APPROACH: BASELINE FIRING RATE EXAMPLE

A limitation of the value coding model presented in Eqs. (1) and (2) is that the firing rate gain and intercept terms *β*_*1*_ and *β*_0_ are degenerate when the learning timescale *τ* is very long. This can make it difficult to determine whether a particular neuron with a long estimated learning timescale is activated by value (*β*_1_ *>* 0) and has a low baseline firing rate, is inhibited by value (*β*_1_ *<* 0) and has a high baseline firing rate, or even whether it is insensitive to value (*β*_1_≈0) and has a moderate baseline firing rate purely based on the data likelihood. To see why, note from Eq. (3) that lim_*τ*→∞_ *v*_*t*_ = *v*_0_, which implies that

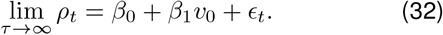

Since there are infinitely many combinations of *β*_0_ and *β*_1_ that satisfy the above equation, the likelihood of the firing rate given the slope and intercept *p*(*ρ*_*t*_ *β*_0_, *β*_1_) is not sufficient to identify the values of these parameters. This means that we need additional information beyond what is available in the data in order to estimate *β*_0_ and *β*_1_. Fortunately, previous work has already established the normal range of baseline firing rates in serotonin neurons. Incorporating this knowledge into our model in the form of a prior distribution for *β*_0_ makes it possible to estimate the values of the remaining parameters, even if the data likelihood is insufficient.

## B PRIOR FOR FIRING RATE INTERCEPTS

### Background

To establish a prior for the firing rate intercepts *β*_0_ in our analysis, we took advantage of previous experiments reported in ref. 9. These experiments are similar to the dynamic Pavlovian experiment analyzed in the main text in important respects:

- Both sets of data consist of *in vivo* tetrode recordings of identified dorsal raphe serotonin neurons in C57BL/6J mice.
- Both sets of data are from head-fixed trace conditioning experiments.

The main differences between the two sets of experiments are as follows:

- The experiments from ref. 9 include reward and punishment conditions, while the dynamic Pavlovian experiment includes only rewards.
- Reward and punishment block transitions were cued in ref. 9, while changes in reward probability in the dynamic Pavlovian experiment were not cued.
- Unconditioned stimuli were deterministic in ref. 9 and stochastic in the dynamic Pavlovian experiment.

In view of these similarities and differences, we took the average baseline firing rate across reward 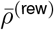 and punishment 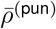 conditions from ref. 9 to be analogous to the baseline firing rate in our model *β*_0_

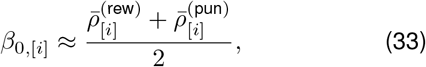

allowing us to estimate the distribution of the firing rate intercept *β*_0_ across neurons from the 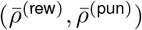 pairs reported in ref. Figs. 3C and 6D.

### Data extraction and pooling

Using Web Plot Digitizer (https://apps.autometris.io/wpd/), we extracted measurements of the mean pre-trial firing rates of *N* = 29 neurons during water reward and air puff punishment blocks of the trace conditioning experiment reported in ref. Fig. 3C, and *N* = 21 neurons during water or chocolate milk reward and quinine solution punishment blocks of the trace conditioning experiment reported in ref. Fig. 6D. Data were pooled across the two experiments because we did not observe significant differences in the distributions of firing rates across the two experiments (differences in 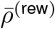 are symmetric about zero, Mann–Whitney U-test *p* = 0.74, differences in 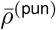 are symmetric about zero, Mann–Whitney U-test *p* = 0.68).

### Results

The distribution of firing rate intercepts implied by ref. 9 is approximately gamma with mean 3.41 Hz and SD 2.34 Hz (*β*_0_ ~ Gamma[2.12, 1.61] using the shape *α* scale *θ* parameterization; maximum likelihood estimates obtained with <monospace>scipy.stats.gamma.fit</monospace>; Fig. S6A). Since our goal is to arrive at a prior distribution for the firing rate intercepts in a new sample of neurons, we quantified the sampling-related uncertainty in our estimated intercept distribution using bootstrapping (Fig. S6B). We found that the bootstrap distribution of the natural log shape and scale parameters is a close match to a multivariate normal

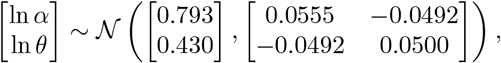

as shown in Fig. S6C, which we therefore used as a prior for the distribution of firing rate intercepts in our analysis.

### Verification

The analogy between the cross-condition average firing rate in ref. 9 and the firing rate intercept *β*_0_ given in Eq. (33) depends on the assumption that the effects of rewards and punishments on the firing rates of serotonin neurons are of roughly equal magnitude and opposite sign, such that they cancel out when averaged. To guard against the possibility that this assumption is wrong, we reserved the baseline firing rates reported in ref. 9 Fig. 8S2A (measured under unspecified conditions) as a test dataset to check the baseline firing rate distribution implied by ref. Figs. 3C and 6D. The distribution of firing rate intercepts implied by Eq. (33) shown in Fig. S6 closely agrees with the distribution implied by the test data (Fig. S7), validating our approach.

**Figure S6.**
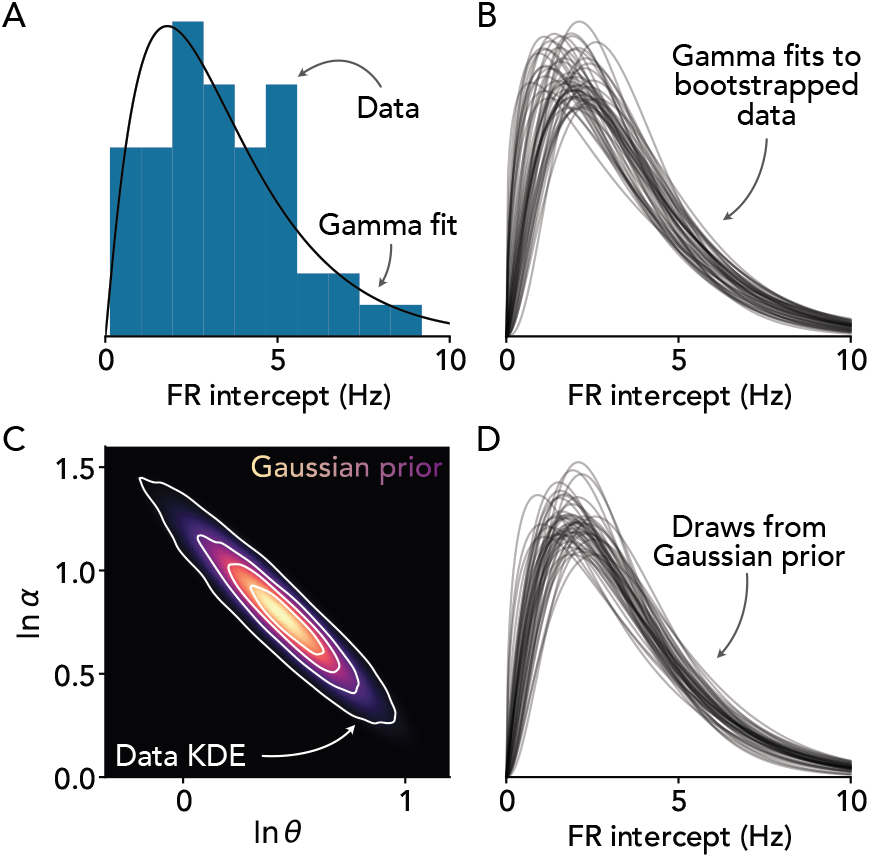
Firing rate intercept prior. **A** Histogram of estimated firing rate intercepts *β*_0_ from ref. 9 with fitted gamma distribution. *N* = 50 neurons from ref. Figs. 3C and 6D. **B** Gamma distributions fitted to bootstrapped firing rate intercept data from A. Note variability due to sampling error. **C** Estimated distribution of gamma distribution parameters across bootstrap samples. Contours show a kernel density estimate of the joint distribution of ln *α* (natural log shape) and ln *θ* (natural log scale) parameters across bootstrap samples. Heatmap shows fitted multivariate Gaussian distribution to be used as a prior. **D** Draws from Gaussian prior distribution over ln *α* and ln *θ*. Note close resemblance to the bootstrap distribution in B.

**Figure S7.**
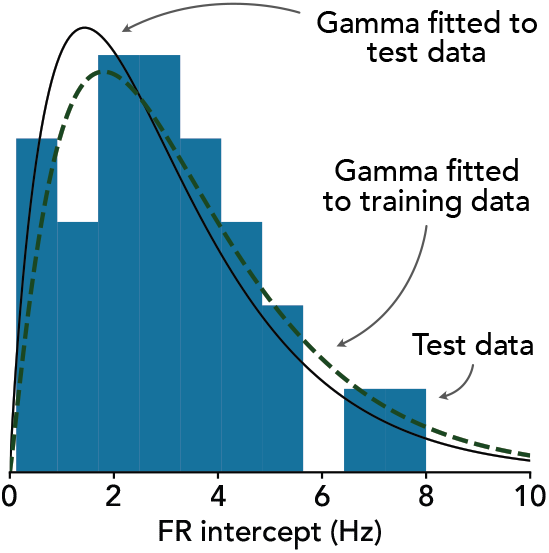
Verification of firing rate intercept prior distribution. Histogram of firing rate intercepts from ref. 9 Fig. 8S2A (test data; *N* = 28 neurons) and fitted gamma distribution (black line) compared with the distribution fitted to data from ref. Figs. 3C and 6D (dashed green line; *N* = 50 neurons).

**Figure S8.**
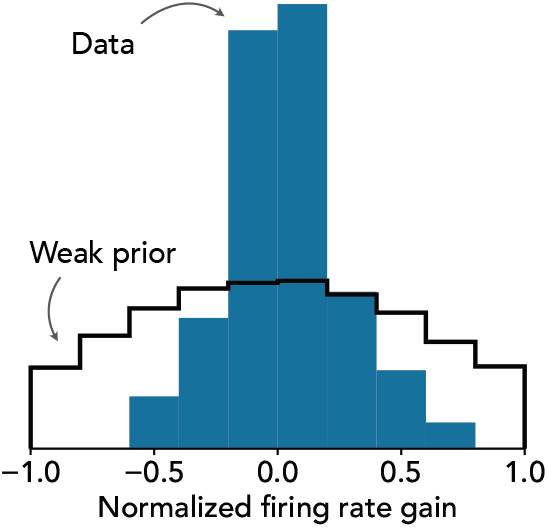
Prior for normalized firing rate gain. Histogram of normalized firing rate gains *β*_1_^*′*^ calculated according to Eq. (34) using data from ref. 9 Figs. 3C and 6D compared with truncated *N* (0, 0.75^2^) prior.

## c PRIOR FOR FIRING RATE GAIN

Using the same data as in Supplementary Note B, we established a prior for the normalized firing rate gain 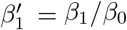 using

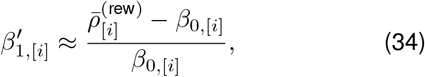

where *β*_0,[*i*]_ is estimated according to Eq. (33) and where we use 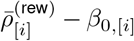 as an estimate of *β*_1_. This estimate is sensitive to mis-estimation of *β*_1_, and, unlike with *β*_0_ (see Supplementary Note B), we are unaware of data that could be used to check our estimate. We therefore use Eq. (34) to set only a weak prior of *β*_1_^*′*^ *~ N* (0, 0.75^2^) truncated to [*⨪*1, 1], as shown in Fig. S8.

## D ALTERNATIVES TO MIXTURE MODELLING APPROACH

Here we have chosen to infer the properties of value, thirst, and null coding serotonin neurons using a mixture model due to important limitations of alternative approaches. In this section, we briefly outline the alternatives considered and why they were not chosen.

### Problem definition

The goals of this work are 1) to measure the proportion of serotonin neurons that encode value and 2) to infer the properties of value coding neurons. To accomplish these goals, we have analyzed a dataset of measurements of the activity of a number of serotonin neurons. The challenges we faced are that 1) the only information we can use to determine whether an individual serotonin neuron encodes value is its activity over time, 2) the time course of activity in a serotonin neuron that encodes value is not necessarily very different than the time course of activity in a serotonin neuron that encodes thirst or where the activity drifts randomly (null coding), and 3) value, thirst, and null coding are particularly difficult to tell apart if the firing rate gain is small and/or if the learning timescale associated with value coding is long. To address these challenges, we needed to select an analysis approach that can cope with unlabeled data (challenge 1), high noise (challenge 2), and confounding that is uneven throughout the parameter space (challenge 3).

### Statistical null hypothesis testing

The most familiar way to address the first challenge listed above would be to use statistical null hypothesis testing to rule out thirst and null coding in a subset of neurons. For example, to rule out null coding, we might have adopted the circular trial permutation test from our previous work (12) and selected only the subset of neurons with activity patterns that are very unlikely under null coding (e.g. at the *p <* 0.05 level) as being non-null-coding. A similar procedure could have been repeated for thirst coding to obtain a set of non-null-, non-thirst-coding neurons. By process of elimination, we would then consider these neurons to be value coding and analyze them further. The basic pattern of this approach will be familiar to most readers: apply a statistical test, label the neurons with *p <* 0.05 as having the coding feature of interest, and then analyze only that subset.

While this approach adequately addresses the first challenge posed above, it does not address challenges two or three. Specifically, because it is generally difficult to tell apart serotonin neurons with value, thirst, or null coding features, and because typical statistical tests focus on limiting the rate of false positives rather than false negatives (including the circular trial permutation test from our previous work, ref. 17), it is likely that a meaningful fraction of value coding serotonin neurons would be discarded. Thus, a bias towards false negatives due to low statistical power could cause the fraction of value coding neurons to be meaningfully underestimated. A further problem with this approach is that the subset of serotonin neurons that *are* selected as value coding will be the neurons with value coding features that are the most different from the activity patterns observed in thirst and/or null coding neurons. In other words, there will be a bias towards selecting serotonin neurons with unusually high firing rate gains and short learning timescales. This phenomenon, whereby selecting data points according to statistical significance causes model parameters to be biased towards extremes, is well documented in the statistical literature (47–49). Overall, taking this approach would likely have led us to conclude that value coding serotonin neurons are less abundant and encode value more strongly and over a shorter timescale than is actually the case.

### Model comparison

A more sophisticated way to address the first challenge listed above would be to fit a value, thirst, and null coding model to each cell, compare their predictive accuracy (using cross-validation, AIC, BIC, etc.), and then label each cell as following the coding scheme corresponding to the best fitting model. Compared with the null hypothesis testing approach outlined above, this approach does not have a strong bias towards thirst or null coding, eliminating the tendency to conclude that value coding neurons are rarer and respond more strongly to recent rewards than is actually the case.

While the model comparison approach represents an improvement over null hypothesis testing in terms of our two main goals, it does not fully address challenges 2 and 3. Specifically, because this approach involves labeling individual neurons as being value, thirst, or null coding in order to conduct further analysis, information about the uncertainty of these labels is lost. This could pose a problem if the sample of serotonin neurons in the dataset analyzed here happens to be enriched in neurons with activity patterns that are only slightly more consistent with value coding than thirst coding, for example, simply due to chance. Since the model comparison approach does not take the strength of evidence for one coding scheme over another into consideration, we would be at risk of confidently concluding that a large fraction of neurons encode value with a low firing rate gain and long learning timescale. The general problem of label-based methods ignoring uncertainty in data labels is briefly discussed in ref. 50.

## Notes

### Competing Interest Statement

The authors have declared no competing interest.

### Summary of Updates

Updated discussion with a new paragraph.

### Funding

- Canadian Institutes of Health Research: PJT- 175319, PJT-175325

